# Model-based differential sequencing analysis

**DOI:** 10.1101/2023.03.29.534803

**Authors:** Akosua Busia, Jennifer Listgarten

## Abstract

Characterizing differences in biological sequences between two conditions using high-throughput sequencing data is a prevalent problem wherein we seek to (i) quantify how sequence abundances change between conditions, and (ii) build predictive models to estimate such differences for unobserved sequences. A key shortcoming of current approaches is their extremely limited ability to share information across related but non-identical reads. Consequently, they cannot make effective use of sequencing data, nor can they be directly applied in many settings of interest. We introduce *model-based enrichment* (MBE) to overcome this shortcoming. MBE is based on sound theoretical principles, is easy to implement, and can trivially make use of advances in modern-day machine learning classification architectures or related innovations. We extensively evaluate MBE empirically, both in simulation and on real data. Overall, we find that our new approach improves accuracy compared to current ways of performing such differential analyses.

---

Using next-generation sequencing, we can now assay up to billions of DNA or RNA sequences in parallel for an ever-expanding set of properties of interest [1– 3]. As a consequence, high-throughput sequencing has dramatically changed the landscape of biological discovery—both for basic scientific inquiry into cellular transcriptomes [4] and protein behavior and evolution [3, 5], and in application areas spanning human disease and variant detection [5, 6], engineering anti-viral immunogens and therapeutics [3, 7, 8], drug and antibiotic resistance [3, 5], regulatory element engineering in synthetic biology [9] and beyond. Across many of these scientific areas, a key desired outcome from a high-throughput sequencing experiment is to quantify the change in relative abundance of a particular sequence between two conditions for a large number of distinct sequences. This type of quantification is often referred to as estimating the “log-enrichment” of a sequence between conditions [2, 5, 7, 8, 10–12]. For example, log-enrichment estimation is performed in differential analyses of RNA-seq and ATAC-seq experiments [4, 13–15] and in high-throughput selection experiments [7, 8, 16–20]. The latter are frequently used for directed evolution [21, 22], deep mutational scanning [2, 3, 5, 10, 12, 23], and functional enrichment analysis [24]. The wide-ranging biologically-significant applications of such selection experiments include: antibody design [25, 26]; profiling pathogen proteomes for epitopes and major histocompatibility complex binding [17, 18]; improving thermostability [27]; assessing binding [12, 16, 21], catalytic activity [27, 28], and packaging efficiency or infectivity of viral vectors [7, 8, 19, 20].

By accurately estimating log-enrichment for large sequence libraries in these settings, one can identify sequences that are more (or less) likely to have desired properties. Consequently, such estimates also have the potential to reveal insights into the sequence determinants of the property of interest. Increasingly, log-enrichment estimates are also being used as supervised labels for training machine learning models so that one may predict enrichment for unobserved sequences, or probe the model to gain further insights [16, 19– 21, 29–32]. These supervised models are often more accurate than popular physics-based and unsupervised machine learning methods such as Rosetta and DeepSequence [32].

## Limitations of log-enrichment estimates

Although standard count-based log-enrichment (cLE) estimates calculated from observed read counts have proven incredibly useful, they suffer from a fundamental shortcoming—the inability to share information across non-identical reads. This inability causes a loss of important available information in a number of practical settings, including:

1. *Short reads*: when short, possibly overlapping, reads are available that each only cover a portion of the *sequence of interest*—*i. e*., the entire span of sequence which we would like to quantify—rather than long reads that span the entire sequence of interest.
2. *Sparse reads*: when few sequencing reads are available per library sequence, as is especially common with long-read sequencing [6, 33–35].
3. *Hybrid reads*: when a combination of long and short reads are collected.
4. *Negative selection*: when the goal is to discover sequences enriched in a property that is opposite from the selection.
5. *More than two conditions*: when we seek to characterize sequences across multiple conditions/selections, such as might occur when engineering gene therapy viral vectors to selectively infect one cell type but not another.

For the case of sparse reads and negative selection (*i. e*., low sequencing counts), it is well-known that cLE estimates suffer from high variance [5, 11, 20]. Previous efforts to reduce variance employ regression to “denoise” cLE estimates by either using a model to intelligently aggregate data across iterative selection rounds [5], or by downweighting examples with low counts [16, 20]. While these techniques can yield higher-quality results, they are extremely limited in their ability to share information across non-identical reads. As a simple, intuitive example, if only 10 out of 300 positions in a sequence of interest are predictive of the property of interest, better statistical power could be achieved by calculating cLE using counts defined only by the 10 relevant positions rather than all 300, since the latter will cause most reads to appear to be non-identical and hence treated separately. A method that could automatically deduce such concepts would, therefore, be of high value. This simple, intuitive idea can be generalized well beyond this example, as we discuss next.

Ideally, to accurately estimate log-enrichment (LE), we would like sequencing data with high read coverage that is comprised of reads that each individually cover the full sequence of interest. However, in practice, individual reads often do not cover the entire sequence of interest—this typically arises with short-read sequencing, but could also occur when using long-read technologies to analyze large sequences of interest [35]. In these settings, it is not obvious how to count reads for the sequence of interest, nor how to calculate the desired cLE estimates. A similar issue occurs in the hybrid read scenario of having both short and long reads. To tackle the LE estimation problem nonetheless, one might consider estimating read-level cLE estimates and then devising a heuristic to add them together to produce a LE estimate for the sequence of interest. However, such an approach is not likely to account for correlations across reads (*e. g*., linkage disequilibrium), nor allow for data sharing by way of partial overlap between reads. Moreover, of the abundance of possible heuristics one might consider, is not clear which to use, and the answer likely depends on the specific application. In applications where there is a known reference sequence—such as in many RNA-seq and ATAC-seq experiments—the reference can help provide information about how to combine reads [4]. However, this is typically performed by alignment and assembly, followed by cLE estimation; thus such approaches also suffer from many of the same limitations just described. Devising an alternative approach to LE estimation—one that is capable of both “sewing” together partially overlapping reads and, more generally, sharing across non-identical reads—would enable more efficient sharing of information.

## A new approach for log-enrichment estimation

Ultimately, a method that can automatically learn to share information as appropriate across non-identical reads will improve our ability to extract important information from sequencing data in a range of settings. Herein, we propose and evaluate a novel, coherent framework that enables us to do just that. Our manuscript is organized as follows: next, we (i) detail how log-enrichment estimates are currently computed; (ii) provide a high-level overview of our approach, *model-based enrichment* (MBE); (iii) provide a detailed empirical characterization of MBE using data from simulated highthroughput selection experiments; and (iv) do the same on real experimental data.

Overall, we find empirically that MBE enables effective analysis across a broader range of common experimental setups than can currently be achieved, including when short-read, long-read, or both types of sequencing reads are used. Our primary motivation is to improve *predictions* of log-enrichment on new (unobserved) sequences, as this is most relevant to our own work in machine learning-guided library design. However, our results show that MBE also enables better *estimation* of log-enrichment, the more classical use case. We show that, compared to existing approaches based on cLE over a broad range of settings, MBE produces predictions that correlate better with true labels. We show that this is, in part, a downstream consequence of the fact that MBE is more robust to low sequencing counts. We also show that MBE enables better characterization of sequences of interest from a negative selection experiment and, as a consequence, is better at identifying sequences that are high in one property while simultaneously low in another—such as we might seek when designing gene therapy viral vectors to infect one cell type and not others—which we call *selectivity* experiments.

## Results

### Overview of current log-enrichment estimation approaches

Given sequencing read counts from two libraries corresponding to two conditions, *A* and *B*, cLE estimates are typically calculated by: (i) indexing the unique sequences in the sequencing data from libraries *A* and *B*, denoting each by *x*_*i*_; (ii) computing read counts, 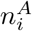 and 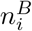, specifying the number of times *x*_*i*_ appeared in the sequencing data from each library; (iii) normalizing the counts 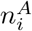 and 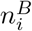 by the total number of reads from each library, *N*^*A*^ and *N*^*B*^; (iv) and, finally, taking the log-ratio of the normalized counts. Thus, the cLE estimate, log *e*_*i*_, is given by: 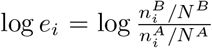. The estimate log *e*_*i*_ has higher variance when the counts 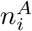 and 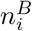 are lower. For example, for fixed *N*^*A*^ and *N*^*B*^, a sequence with 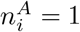 and 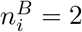 has the same log *e*_*i*_ as a sequence with 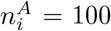 and 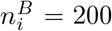, yet the latter is supported by 100 times more evidence (and thus is a lower-variance estimate).

Supervised machine learning regression models have been used to reduce the variance of (*i. e*., “de-noise”) such cLE estimates [5, 16, 20], and to make predictions for sequences not present in the training library [19–21, 32]. The latter strategies, which we refer to as *LE regression* approaches, use cLE estimates as supervised labels to learn a predictive model mapping from sequence to predicted LE. Zhu *et al*. [20] additionally derive a variance estimate for log *e*_*i*_ which enables them to weight each training sequence according to the amount of evidence that supports its cLE estimate, yielding improved predictive performance. Consequently, when comparing to a baseline for LE prediction, we use this approach, which we refer to as *weighted LE regression* (wLER).

### A new approach: model-based enrichment

Regression-based LE estimation (or prediction) is performed in two sequential steps: first, one computes a cLE estimate for each unique sequence [2, 5, 10, 11], and second, one trains a regression model to predict these cLE estimates from the observed sequences, possibly weighting each sequence to account for its corresponding level of evidence [20, 32].

We introduce a new method, MBE, that performs both of these steps at once, resulting in a more powerful and more general analysis framework. We do so by reframing the LE estimation problem: we show that a cLE estimate can be viewed as an approximation to the log of what is known as a *density ratio*—the ratio of probability densities of the observed sequence under each condition. Therefore, we can estimate and predict LE by solving a *density ratio estimation* problem (DRE). Further, DRE can be effectively and accurately performed by training a probabilistic classifier to predict which of the two densities a sample came from (*e. g*., condition *A* or *B*) [36–42]. Specifically, the ratio of such a classifier’s predicted class probabilities provably converges to the density ratio [37–39].

Through this series of theoretically-justified steps, we are able to transform the problem of estimating LE into one of training a read-level classifier to distinguish which condition a read came from. This transformation provides several distinct advantages over existing methods, which we outline here and, later, demonstrate empirically. The first advantage is that we can readily make use of modern-day neural network models in a plug-and-play manner, which also enables us to easily handle reads (possibly overlapping) of different lengths. For example, fully convolutional neural network classifiers naturally handle variable-length sequences because the convolutional kernels and pooling operations in each layer are applied in the same manner across the input sequence, regardless of its length. Similarly, the same can be said for transformer architectures. Moreover, because MBE is classifier-based, it is easy to implement using standard software packages: one need simply train a classifier using any standard classification tools. A second advantage is that our approach naturally accounts for differing levels of evidence per sequence of interest, which in previous LE regression methods was either ignored or addressed *post hoc* under specific distributional assumptions [16, 20]. Third, our approach trivially generalizes to settings with more than two conditions of interest by replacing the binary classifier with a multi-class one; this enables us to naturally handle experiments with multiple rounds of selection or properties of interest.

We highlight that our classifier-based DRE approach differs substantially from several recent approaches that also make use of classification. In one, cLE estimates are thresholded and a classifier built to predict the resulting binarized labels (*e. g*., [19]). In another, a classifier is built to predict whether a sequence appeared at all in post-selection sequencing data (*e. g*., [30]). Neither of these approaches address the shortcomings that we seek to resolve with MBE.

#### Technical overview of MBE

Here, we provide more detail about our MBE approach (see Methods for full detail). Recall that the cLE estimate is the log-ratio of the two normalized counts, 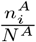 and 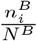. These normalized counts are also the empirical frequencies of the *i*^th^ unique read, *x*_*i*_, in the sequencing data for conditions *A* and *B*, respectively. In particular, these two ratios are the sample-based estimates of the population frequencies of *x*_*i*_ in each library. We denote the population frequencies by the probabilities *p*^*A*^(*x*_*i*_) and *p*^*B*^(*x*_*i*_). Consequently, log *e*_*i*_ can be viewed as a sample-based estimate of the population-level LE, which we denote log *d*(*x*_*i*_). Specifically, 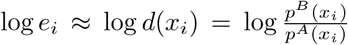, where *d* is the *density ratio* between the library distributions. By training a binary classifier with parameters, *θ*, to predict the probability that a read with sequence *x*_*i*_ came from library *B, p*_*θ*_(*l* = *B* | *x*_*i*_), we can estimate log *d*(*x*_*i*_), and hence the LE, as 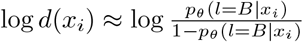 [37, 39]. It has been proven theoretically that under a correctly specified model, this density ratio estimation method is optimal among a broad class of semi-parametric estimators—that includes the wLER method—in terms of asymptotic variance [37] (Supplementary Note).

### Overview of experimental setup

Next, we describe the simulated and experimental datasets used to empirically compare and contrast our MBE approach with cLE and wLER across a broad range of settings. Then, we provide an overview of the evaluation metrics, before finally presenting the simulated and experimental results.

#### Simulated data

Using simulated experiments, we sought to understand the strengths and weaknesses of the MBE and wLER approaches as we changed following simulation settings:

1. the length of the sequence of interest, *L*, ranging from 21–2,253 nucleotides.
2. whether short or long reads were used (300 *vs*. 10,000 nucleotides).
3. the number of unique sequences in the theoretical pre- and post-selection libraries, *M*′, ranging from 8.5 × 10^6^–2.6 × 10^7^.
4. the number of pre- and post-selection reads, *N*^pre^ and *N*^post^—always set equal to each other, ranging from 4.6 × 10^3^–4.6 × 10^7^.
5. the complexity of the functional mapping between sequence and property of interest; this complexity was characterized in terms of a summary parameter controlling the amount of epistasis, *T*. We simulated libraries that correspond to three types of experimental library constructions:
  a. *Insertion* of a sequence into a fixed background. In a given library, the insertion has fixed-length and a fixed position within the background sequence. The insertion library construction is motivated by our work in adeno-associated virus (AAV) capsid engineering which aims to understand sequence determinants of AAV properties such as packaging [20]. In this study, the sequence of interest is a 21-mer nucleotide insertion sequence into the capsid with fixed background. Herein, we simulate this insertion library with varying lengths (21, 150, and 300 nucleotides). The pre-selection library is generated to be roughly uniform in nucleotide space (technically, an “NNK” degenerate codon distribution).
  b. *Random mutagenesis*—motivated by a study to understand the fitness landscape of a green fluorescent protein of length 714 nucleotides [31]. Herein, we mutagenise the green fluorescent protein across all positions using a 10% mutation rate to generate the pre-selection library.
  c. *Recombination*—motivated by an AAV directed evolution study [8], wherein several AAV serotypes are recombined using seven crossovers separating eight recombination blocks. Herein, we generate library sequences by recombining nine AAV serotypes using eight equally-sized blocks. The total length of all eight blocks is 2253 nucleotides.

A summary of the simulated sequencing datasets is provided in Table 1.

**Table 1.** Summary of simulated datasets. For each dataset we list the: library name (Library), sequence length in nucleotides (*L*), number of unique library sequences (*M* ^*′*^), epistasis hyperparameter used for fitness simulation (*T*), read type (short, long, or hybrid), % of the sequence of interest covered by individual reads (Cover), and number of pre-selection and post-selection reads (*N*^pre^ and *N*^post^), which were always equal. We simulate 4.6 *×* 10^7^ short reads to match the experimental data from Zhu *et al*. [20], and up to 4.6 *×* 10^5^ long reads to be within the current throughput of PacBio’s technologies [34, 35]. Each dataset is described in more detail in the Methods.

| Library | $L$ | $M'$ | $T$ | Read Type | Cover | $N^{\text{pre}} = N^{\text{post}}$ |
| --- | --- | --- | --- | --- | --- | --- |
| 21-mer insertion | 21 | $8.5 \times 10^6$ | 140 | Short | 100 | $4.6 \times 10^7$ |
| 150-mer insertion | 150 | $8.5 \times 10^6$ | 1000 | Short | 100 | $4.6 \times 10^7$ |
| 300-mer insertion | 300 | $8.5 \times 10^6$ | 2000 | Short | 100 | $4.6 \times 10^7$ |
| avGFP mutagenesis | 714 | $2.5 \times 10^7$ | 4760 | Long | 100 | $4.6 \times 10^5$ |
| avGFP mutagenesis | 714 | $2.5 \times 10^7$ | 4760 | Short | 42 | $4.6 \times 10^7$ |
| AAV recombination | 2253 | $2.6 \times 10^7$ | 15020 | Long | 100 | $4.6 \times 10^5$ |
| AAV recombination | 2253 | $2.6 \times 10^7$ | 15020 | Long | 100 | $4.6 \times 10^4$ |
| AAV recombination | 2253 | $2.6 \times 10^7$ | 15020 | Long | 100 | $4.6 \times 10^3$ |
| AAV recombination | 2253 | $2.6 \times 10^7$ | 15020 | Short | 13 | $4.6 \times 10^7$ |
| AAV recombination | 2253 | $2.6 \times 10^7$ | 15020 | Hybrid | 100 long<br>+<br>13 short | $4.6 \times 10^3$ long<br>+<br>$4.5 \times 10^7$ short |

Underlying each of the motivating selection experiments is a property on which the sequences get selected, such as protein fluorescence. To simulate selection, we must simulate the ground truth *fitness function* that maps sequence to property. We did so as a linear function of a number of features, including: all independent amino acid sites, and *T* higher-order epistatic features drawn randomly from all possible such effects, in a manner that recapitulates the distribution of these effects in a real protein fitness landscape. In particular, combining insights from several papers [43–45], we assumed that *T* scaled linearly with the length of the sequence of interest, with a fixed coefficient based on Poelwijk *et al*. [29].

Finally, the process to simulate reads from the pre- and post-selection libraries can be summarized as follows: first, we generate library sequences using one of the three previously described library construction simulations. Then, we randomly perturb the empirical distribution of the simulated library sequences (which simulates slight distributional perturbations that may occur with PCR amplification) to generate a pre-selection probability distribution. Next, the corresponding post-selection probability distribution is determined by scaling the pre-selection distribution according to the simulated fitness of the library sequences. Then, we sample reads that cover the full sequence of interest from the pre- and post-selection distributions. When simulating short reads, we truncate each of these reads to 300 nucleotides, at a position chosen uniformly at random.

We also perform negative selection simulations, which were motivated by experiments wherein one seeks to identify sequences with a property, such as low-binding affinity, for which the only available assay enriches for the opposite, such as high-binding. This situation arises, for example, in studies of AAV tropism [8, 46] where the ideal viral vector selectively infects one cell type, but not others. We, therefore, aimed to estimate the accuracy of wLER and MBE to negatively select against an undesirable fitness and, moreover, to identify sequences of interest that are *selective*—meaning that they are simultaneously high in one fitness (the *positive fitness*) and low in a second (the *negative fitness*). To do so, we simulated two independent fitness functions and used each, separately, on the same pre-selection library to simulate two post-selection libraries and corresponding reads.

Although most of our simulations did not include sequencing errors, we constructed versions of two of the aforementioned datasets that did. For one of the insertion datasets, we used a uniform random substitution error rate of 1%, consistent with observed error rates of Illumina’s next-generation sequencers [47]. For one of the recombination datasets, we used SimLoRD [48] to simulate PacBio SMRT sequencing errors. As shall be seen, the noise had little effect on our results.

### Real experimental data

We also used five experimental datasets—each comprised of sequencing data from a pre-selection library and after one or more selections on that library. For our evaluations, we also used low-throughput experimental property measurements corresponding to the selected property for each of the five sequencing datasets. Each experimental dataset and its corresponding property measurements are summarized in Table 2 and described briefly in the same order here:

**Table 2.** Summary of experimental datasets. For each dataset we list the: library description (Library); reference (Ref.); sequence length in nucleotides (*L*); number of unique library sequences after holding out experimentally-validated variants, if needed (*M*′); number of experimentally-validated variants (*n*); % of the sequence of interest covered by individual reads (Cover); number of pre-selection reads (*N*^pre^); and number of post-selection reads (*N*^post^). For the dataset from Huisman *et al*. [17], the number of reads for each round of selection is presented on a separate row.

| Library | $L$ | $M'$ | $n$ | Cover | $N^{\text{pre}}$ | $N^{\text{post}}$ |
| --- | --- | --- | --- | --- | --- | --- |
| AAV5 insertion [20] | 21 | 8,552,729 | 5 | 100 | 46,049,235 | 45,306,265 |
| SARS-CoV-2-derived peptide [17] | 45 | 167,841 | 24 | 100 | 44,073 | 88,032 |
| SARS-CoV-2-derived peptide [17] | 45 | 167,841 | 24 | 100 | 88,032 | 169,730 |
| SARS-CoV-2-derived peptide [17] | 45 | 167,841 | 24 | 100 | 169,730 | 235,787 |
| SARS-CoV-2-derived peptide [17] | 45 | 167,841 | 24 | 100 | 235,787 | 160,863 |
| GB1 double site saturation [12] | 168 | 536,953 | 11 | 100 | 324,434,913 | 262,112,210 |
| Chorismate mutase homolog [28] | 288 | 3,063 | 11 | 100 | 1,228,687 | 1,929,212 |
| Bgl3 random mutagenesis [27] | 1506 | 468,194 | 6 | 100 | 1,177,842 | 710,555 |

1. A library of 21-mer nucleotide insertions into a fixed AAV background sequence subjected to a round of packaging selection, and packaging titer measurements for five sequences not present in the library [20].
2. A library containing every 15 amino acid peptide in the SARS-CoV-2 proteome (which has 14,439 amino acids) subjected to four rounds of selection for binding to human major histocompatibility complex (MHC). For ground truth, there are *IC*_50_ measurements for 24 peptides [17] held out from the LE analysis.
3. A site saturation mutagenesis library containing all single and double amino acid mutations within the 168 nucleotide IgG-binding domain of protein G (GB1) subjected to selection for binding to IgG-FC. For ground truth, the are Δln(*K*_*A*_) measurements for 11 individual variants held out from the sequencing data [12].
4. A library containing natural chorismate mutase homologs and designed sequences sampled from a direct coupling analysis (DCA) model. All sequences are of length 288 nucleotides. For ground truth there are biochemical measurements for 11 variants held out from the sequencing data [28].
5. A *β*-glucosidase enzyme (Bgl3) error-prone PCR random mutagenesis library subjected to a heat challenge and high-throughput droplet-based microfluidic screening. All sequences are of length 1506 nucleotides. For ground truth, there are *T*_50_ (temperature where half of the protein is inactivated in ten minutes) measurements for six mutants held out from the sequencing data [27].

### Model architectures

We implemented wLER and MBE using several model architectures. To enable direct comparison of the two methods, we kept the set of architectures and allowed hyper-parameters the same for both approaches, excluding the final layer and loss which dictate whether the model is for regression (wLER) or classification (MBE). Specifically, we used the seven model architectures in Zhu *et al*. [20]—three linear models and four fully-connected neural networks (NNs)—as well as four additional convolutional neural network (CNN) architectures. As the linear and NN architectures and hyper-parameters are from a paper that used wLER, to the extent the selected architectures may favor one of the approaches compared herein, they would favor wLER. The CNNs can operate on variable-length sequences, allowing us to train on short reads and make predictions on full-length sequences of interest.

Several of our experiments simulate negative selection against an undesirable fitness, and selectivity experiments that select for sequences that are simultaneously high in a desirable positive fitness and low in an undesirable negative fitness. For simplicity, in these experiments we used only one model architecture—the smallest NN architecture—as a two-output model, one for the positive fitness and one for the negative. We used this architecture because it was the simplest non-linear model architecture we explored—meaning it is capable of capturing higher-order epistasis whilst being relatively parsimonious. Based on the results of our initial simulation experiments, this choice of architecture does not systematically benefit either of the wLER or MBE approaches (Supplementary Fig. 1).

For all real experimental datasets (except for the Bgl3), we similarly used the smallest NN architecture because it tended to achieve better cross-validation performance than the linear architectures and comparable performance to the larger NN and CNN architectures, whilst being more parsimonious (Supplementary Fig. 8a-l). For the Bgl3 dataset, we used a simpler linear model because overfitting was observed with the NNs (Supplementary Fig. 8m-o). For the one dataset that had multiple rounds (Huisman *et al*. [17]), we used a multi-output model with one output per round and took the final prediction to be the average of the predictions for each round.

Note that, in practice, when classes are highly imbalanced (*i. e*., *N*^pre^ is much larger than *N*^post^, or vice versa), the read-level classifier underlying MBE may implicitly learn to predict the more prevalence class, which may be undesirable. To address this, it may be preferable to use standard machine learning techniques for training classification models under class imbalance, such as class weighting wherein samples from each class are weighted so that the classifier learns equally from both classes. Such techniques are not necessary in our simulations because classes are exactly balanced (*i. e*., *N*^pre^ = *N*^post^), but we use class weighting to train read-level classifiers on the experimental sequencing datasets.

### Evaluation methods

For both real and simulated data, we compared and contrasted three approaches, as appropriate: standard cLE, wLER (recall this is a weighted regression on cLE), and our MBE approach which bypasses computation of cLE. For all but cLE, we can both (i) make predictions on sequences not seen in the training data, and (ii) make model predictions on the training data itself to yield LE estimates—a sort of”de-noising” of the cLE estimates. We refer to these two tasks, respectively, as *prediction* and *estimation*. The cLE approach can only be used for estimation, hence it does not appear in prediction experiments.

To compare wLER to MBE on any given dataset, we used all model architectures and hyper-parameters for both methods, and then selected the best combination separately for each of wLER to MBE. No model or hyperparameter selection is required for cLE since it does not use any model or have any parameters.

An important point to appreciate throughout our work is that we cannot use straight-forward cross-validation to assess accuracy because we cannot use the ground truth fitness values to train, only to evaluate. Nor can we use, say, cLE estimates for cross-validation, as these are not ground truth values. Hence in simulated settings, we perform modified cross-validation where we evaluate performance on each fold by comparing predictions to the sequences’ ground truth fitness values. For the real experimental datasets where ground truth fitness values for the library sequences are unknown, we use available low-throughput (non-sequencing-based) experimental fitness measurements (which may still be corrupted by noise, but are more direct measurements of the property of interest than the sequencing-based assays) for validation.

In our simulations, we use three-fold cross-validation to compute the Spearman correlation between ground truth fitness and predicted LE to compare and contrast the performance of each method. Additionally, we make use of a generalized Spearman correlation that focuses on sequences that have the highest ground truth LE—the focusing is controlled by a threshold on true LE which we sweep through a range of values, such that at one extreme, we compute the Spearman of all sequences in the test set, and on the other, of only the most truly enriched sequences (similarly to Zhu et al. [20]). The test set is always comprised of full sequences of interest, even when the training data contained reads that were shorter. For all cross-validation experiments, we averaged the Spearman correlations computed on each fold to produce one cross-validated correlation value. We use William’s t-test to assess statistical significance of the difference between the cross-validated Spearman correlations.

Each selectivity simulation is defined by two different simulated fitnesses, a positive fitness and negative fitness. We learn two-output models (one output per fitness) on these data. We define the selectivity of a sequence as the difference between its positive and negative fitness values. We apply the generalized Spearman correlation evaluation method described above for the positive fitness. For the negative fitness, we use a similar generalized Spearman correlation that focuses on sequences with *lowest* —instead of highest—ground truth LE. In the selectivity experiments, we also seek to compare how well wLER and MBE identify test sequences with high selectivity. To do so, for each method, we rank the sequences in each test fold according to predicted selectivity— the difference between predictions for each fitness—and take the top ten test sequences. Then, we compare the two ground truth fitness values of each of the chosen sequences to the fitness values of a theoretical optimally-selective sequence that has the maximum true positive fitness and minimum true negative fitness observed in the given dataset. We also use McNemar’s test to assess the statistical significance of the difference between the methods’ accuracy at identifying the 1% of test sequences with highest selectivity.

On real experimental data, we compare the wLER and MBE approaches by computing Spearman correlation between predicted LE and low-throughput experimental property measurements. We use a paired t-test to assess statistical significance of the performance difference between wLER and MBE aggregated across all five experimental datasets.

### Results on simulated data

Across all simulated datasets, our MBE approach made significantly more accurate LE predictions than wLER (Fig. 1a) according to standard Spearman correlation (*p* < 10^−10^). The improvements of MBE over wLER in terms of Spearman correlation values were as much as 0.561 and as little as 0.005, with an average of 0.177. In no cases did MBE do worse than wLER. We also found that our MBE method performed better when faced with both Illumina- and PacBio-like sequencing error (Fig. 1, Supplementary Fig. 5). Moreover, MBE was much less sensitive to the choice of model architecture, to such an extent that even the worst-performing MBE model performed better than the best-performing wLER model on several datasets (Supplementary Fig. 1a). Similarly, for the estimation task, MBE outperformed wLER across all simulated datasets (Fig. 1b, Supplementary Fig. 1b). Collectively, these results demonstrate a clear win for MBE over wLER, across a broad range of settings. In the subsequent sections, we examine the following specific settings to get a fuller view of the strengths and weaknesses of each method: sparse reads, overlapping short-reads, hybrid long- and short-reads, negative selection, and selection for sequence selectivity.

**Fig. 1.**
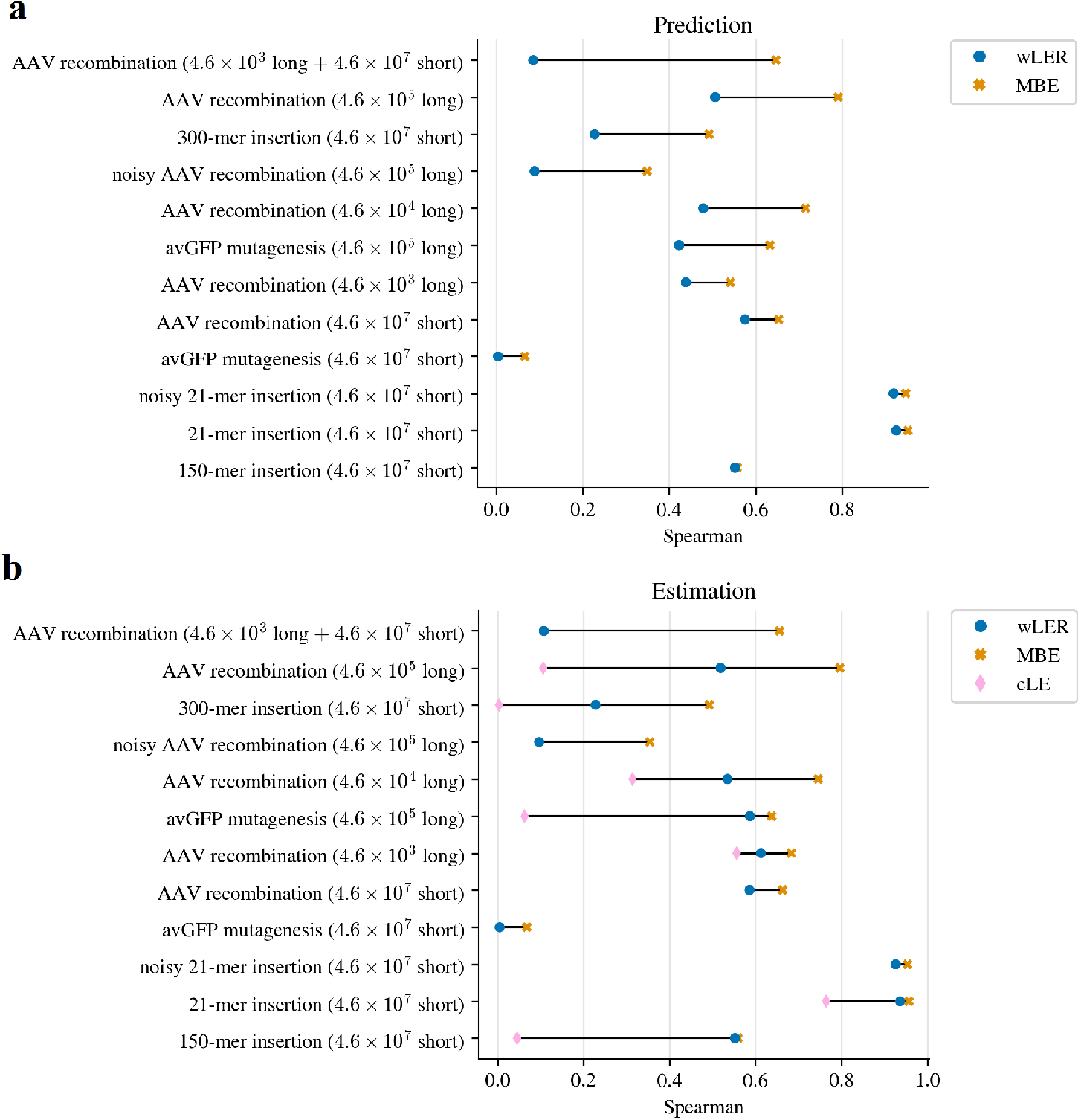
Simulated library results. Spearman correlation between ground truth fitness and cLE, wLER, and MBE estimates on full-length sequences of interest for the tasks of (a) prediction and (b) estimation. For wLER and MBE, each panel displays the Spearman correlation achieved by the best-performing model architecture for each method on each simulated dataset. The cLE approach can only be used for estimation (not prediction), and additionally, only for experiments where the sequencing reads were long enough to cover the sequences of interest (“Cover=100”). Thus, cLE is missing from some experiments. All differences are statistically significant (*p* < 10^−10^). Results shown are for the best architecture for each approach, as described in main text. For comprehensive results across all model architectures see Supplementary Fig. 1.

#### Sparse read setting

We define the sparse read setting as occurring when the average number of sequencing reads per library sequence was lower than 0.02. In our experiments, this includes simulated long-read datasets for the avGFP mutagenesis and AAV recombination libraries. We hypothesized that the MBE approach would have a particular advantage in this setting because of its improved ability to combine information across similar but non-identical reads. On the prediction task, MBE maintains comparable accuracy to wLER on test sequences with high ground truth fitness, while improving accuracy in the other regimes (Supplementary Fig. 2a-b, Supplementary Fig. 3). Additionally, MBE had lower variance than wLER across the different test folds (Supplementary Fig. 3). We also note that the longer the sequence of interest, the more MBE outperforms wLER—this nicely matches our intuition as the longer the read, the more sparse the setting (Fig. 1a, Supplementary Fig. 3d-e, Supplementary Fig. 4). We observed similar trends for the estimation task (Fig. 1b, Supplementary Fig. 1b, Supplementary Fig. 6). When we increase the total number of long reads for the AAV recombination library (from 4.6 × 10^3^ to 4.6 × 10^5^), more unique sequences with low counts occur in the data (Supplementary Fig. 4). Consequently, wLER is particularly challenged because it is trained using cLE estimates that cannot share data across non-identical reads to mitigate the effects of low counts. In fact, wLER is so challenged that, for many model architectures, its performance degrades when provided with more long-read sequencing data (Supplementary Fig. 2a-c). In contrast, MBE follows a more intuitive pattern: more training data always either maintained or improved performance, but never hurt the overall performance metrics (Fig. 1, Supplementary Fig. 2).

#### Short- and hybrid-read settings

In practice, experimenters often offset the sparsity of long-read sequencing by augmenting with higher-throughput short-read sequencing. We refer to this as the hybrid-read setting. Again, our results follow our intuition: for shortread and hybrid datasets, MBE outperformed wLER (Fig. 1a, Supplementary Fig. 2d-f). In fact, because wLER cannot leverage partial overlap between reads, its accuracy actually decreased when long-read data was supplemented with additional short reads, despite the fact that this creates a larger overall training set. In contrast, MBE, again, behaved more intuitively: its accuracy improved with this larger dataset.

#### Negative selection

In negative selection experiments, the property being selected for is opposite from the property of interest. Thus, a key goal is to produce accurate predictions for sequences with low ground truth fitness, for which the post-selection read counts are, by definition, low, making these estimates extremely challenging. We compared wLER and MBE predictive accuracy using generalized Spearman correlation focused on sequences with low ground truth fitness. MBE achieved higher predictive accuracy, not only overall, but also specifically on the subset of the test sequences with lowest true fitness (Fig. 2).

**Fig. 2.**
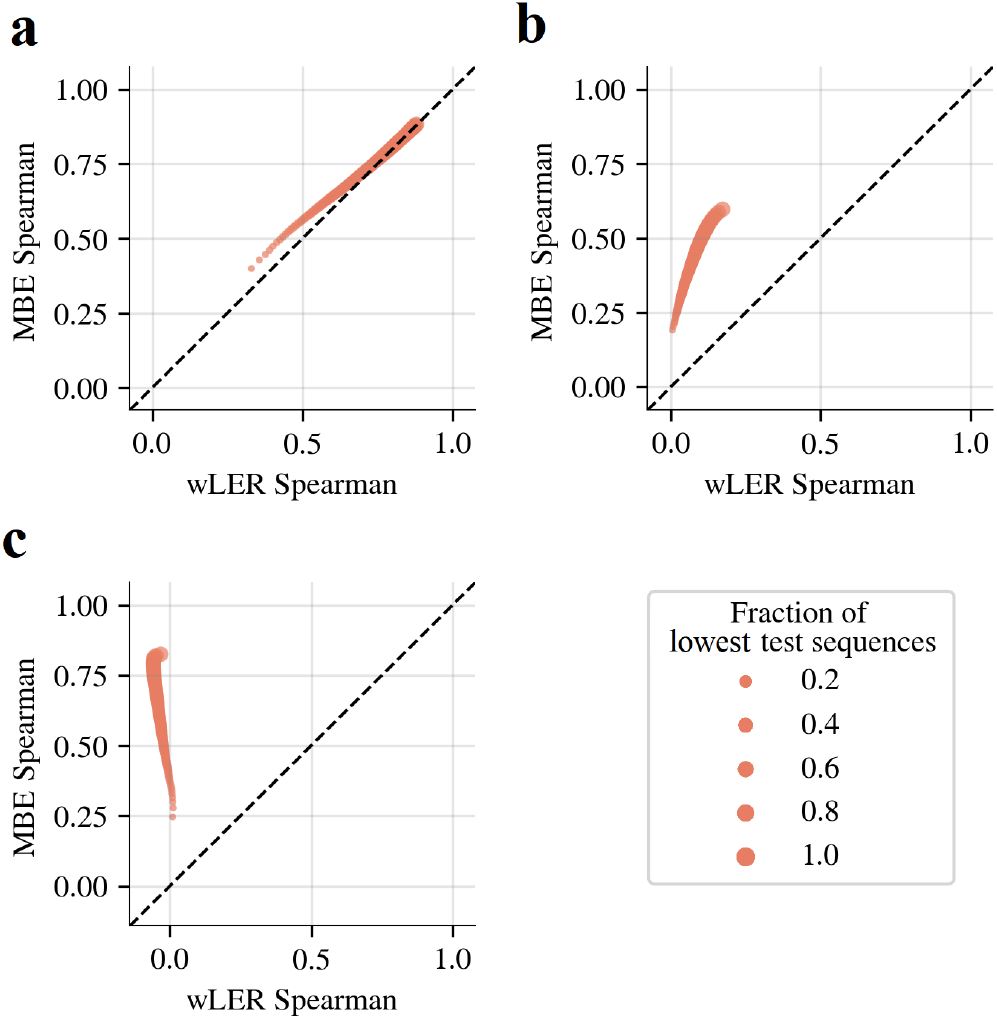
Simulated negative selection prediction results. Comparison of wLER and MBE predictive accuracy for simulated negative selection using the 100-unit NNs on the (a) 21-mer insertion (4.6 × 10^7^ short reads), (b) avGFP mutagenesis (4.6 × 10^5^ long reads), and (c) AAV recombination (4.6 × 10^5^ long reads) datasets. Dot size represents the fraction of test sequences with lowest ground truth fitness used to compute Spearman correlation. In these experiments, we focus on sequences with lower ground truth fitness, which are the smaller dots. The dashed black line represents equal performance of the two approaches.

#### Selection for sequence selectivity

A key reason to seek high predictive accuracy for the negative selection task is so that we can leverage this task to perform a selectivity experiment, wherein we seek to identify sequences that simultaneously appear high in a positive fitness, and low in the negative fitness. Indeed, we find that MBE is better than wLER at identifying those *selective sequences*. First, we found that MBE yielded better predictive accuracy on both fitnesses than wLER (Supplementary Fig. 7a-b, d-e, and g-h). More importantly, MBE was also better at identifying the selective sequences, which we assessed as follows. To measure a sequence’s selectivity, we computed the difference between its positive and negative fitness values: the larger this difference, the more selective the sequence is for the positive selection relative to the negative selection. MBE was more accurate than wLER in identifying selective sequences (Fig. 3). Moreover, the best sequences identified by MBE were, on average, closer to a theoretical optimally-selective sequence, compared to wLER (Fig. 3, Supplementary Fig. 7c, f, and i). Overall, for each of the three datasets, MBE was significantly better than wLER at identifying the 1% of test sequences with highest true selectivity (*p* < 10^−3^).

**Fig. 3.**
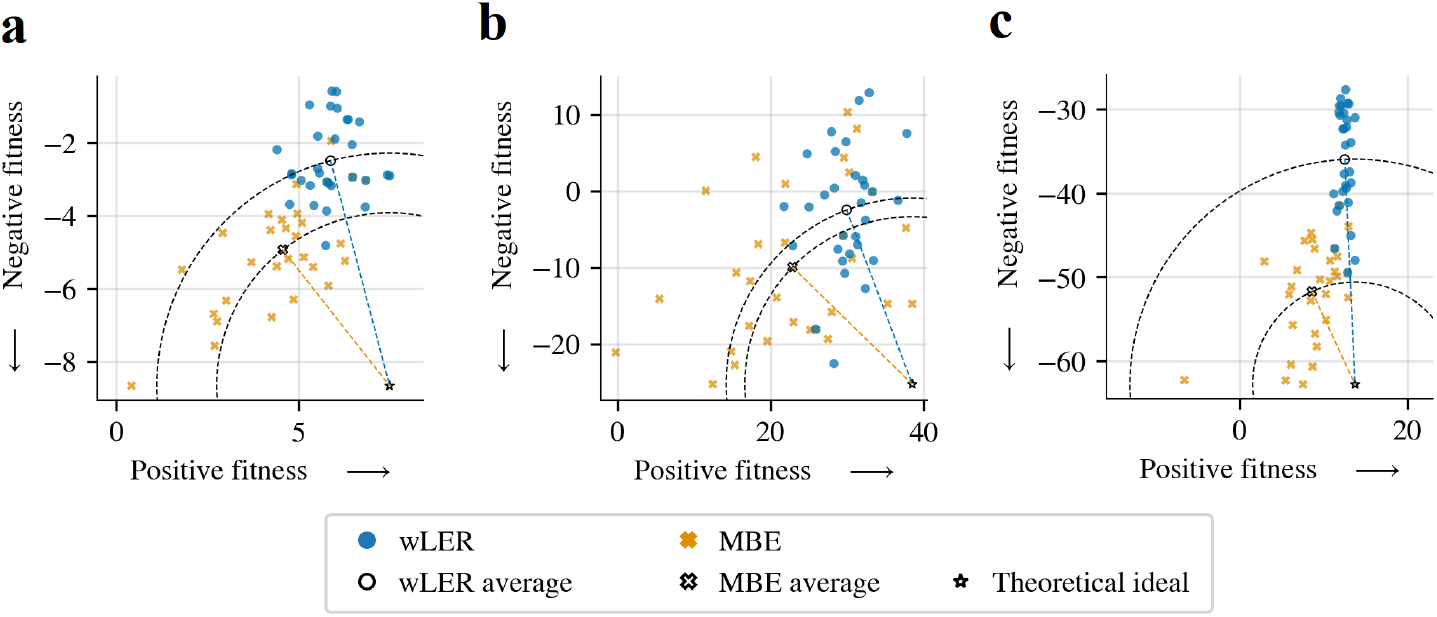
Simulated sequence selectivity prediction results. Comparison of wLER and MBE (using 100-unit NNs) for identifying selective test sequences over three simulated datasets: (a) 21-mer insertion (4.6 × 10^7^ short reads), (b) avGFP mutagenesis (4.6 × 10^5^ long reads), and (c) AAV recombination (4.6 × 10^5^ long reads). Colored points show the true positive and negative fitness of the top ten test sequences identified from each of three test folds from three-fold cross-validation according to each model’s predicted selectivity (*i. e*., difference in predicted positive and negative fitness values). To gauge overall performance, the average point from each method is also plotted in black-and-white, as is a theoretical optimally-selective sequence (star) with the maximum positive fitness and minimum negative fitness among all sequences in the relevant dataset. Distance from optimal to average is conveyed by a circular contour line through the average point for each method; the size of the gap between the two circles is indicative of how much closer MBE is to the optimum than wLER. On all three datasets, MBE is significantly more accurate than wLER at identifying the 1% of test sequences with highest true selectivity (*p* < 10^−3^).

### Results on real experimental data

Having characterized the behavior of wLER and MBE in a broad range of simulated settings, we applied these methods on real experimental data. Across all the real datasets, MBE achieved better predictive accuracy than wLER (Fig. 4, Supplementary Fig. 9). Recall that an important challenge here is that, to obtain the best ground truth experimental values possible, we require access to detailed biophysical assays rather than sequencing-based proxies. Consequently, the validation data we have access to have extremely limited sample sizes (ranging from 5–24 test points), thereby limiting our our ability to detect statistical significance on each dataset individually. Nevertheless, the trends that we observed on the simulated data continue on each dataset, and when performance over all of them is considered jointly, the improvement of MBE over wLER is statistically significant (*p* < 0.03) (Fig. 4).

**Fig. 4.**
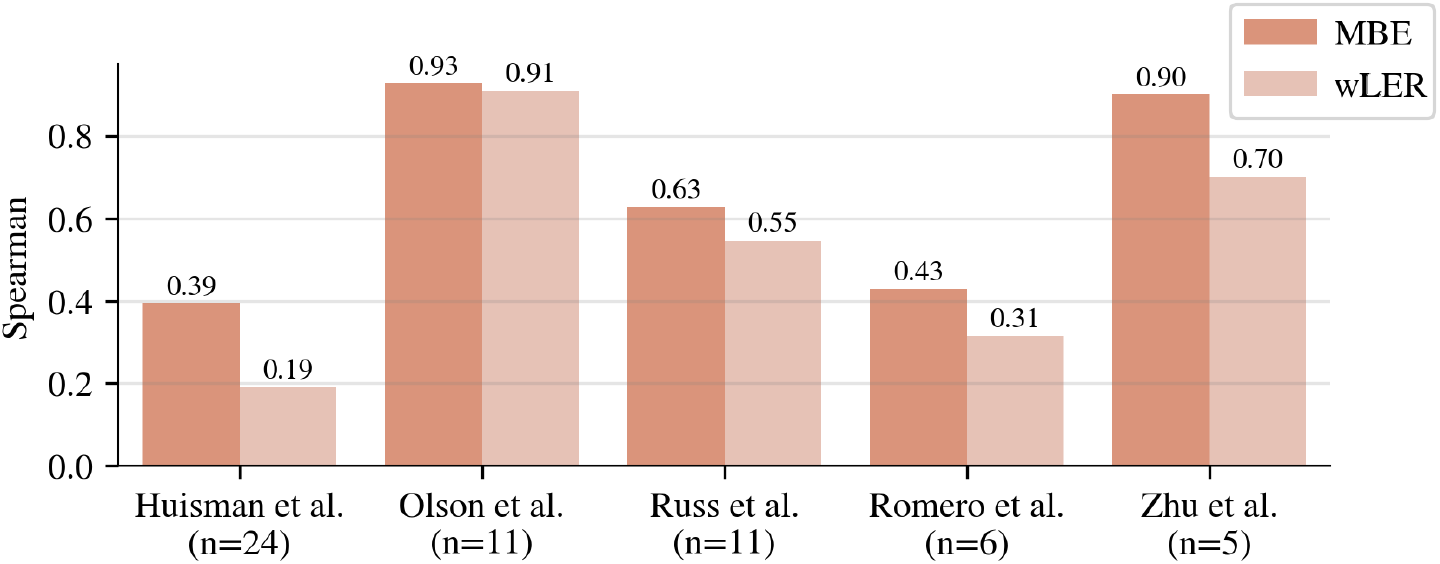
Real experimental prediction results. Comparison of Spearman correlation between wLER or MBE predictions and *n* experimental property measurements from Zhu *et al*.[20], Huisman *et al*.[17], Olson *et al*.[12], Russ *et al*.[28], and Romero *et al*.[27]. Each method is trained on real pre- and post-selection sequencing data, then used to predict the fitness of the *n* unobserved test sequences. The 100-unit NN model architecture is used for all datasets except that from Romero *et al*.[27], for which the linear architecture with IS features is used. The average performance improvement of MBE over wLER over all five experimental datasets, jointly, is statistically significant (*p* < 0.03).

As an aside, for the SARS-CoV-2 dataset from Huisman *et al*. [17], we found that predictions of experimental *IC*_50_ by MBE were more accurate than the predictions by NetMHCIIpan4.0, a model specifically devised to predict peptide binding to MHC II molecules (Supplementary Table 1).

## Discussion

Quantitatively characterizing the difference in sequence abundances between two conditions using high-throughput sequencing data—as occurs, for example, in high-throughput selection experiments—is a key component in answering a large range of scientific questions. Not only do we wish to quantify the differences in observed data, but we also often want to predict the difference for sequences not yet observed, in order to design further rounds of experimentation, for example. Until now, quantification was accomplished by counting the number of times a sequence occurred in each condition and taking the ratio of these counts (after normalization). Then, optionally, one may have constructed a regression model to predict these count-based log-enrichment ratios. The key issue underlying all of this processing is the inability of count-based estimates to share any information across sequences that are not identical, when such sharing of information can be extremely valuable. Herein, we introduce, and evaluate, a framework that overcomes this key limitation. This framework is based on a reformulation of the problem that uses density ratio estimation, implemented using any standard machine learning classifier. Our new method, model-based enrichment, improves performance over competing approaches based on either raw counts or weighted regression on count-based log-enrichment. In particular, we show this improvement holds across a broad range of simulated data, as well as on real experimental data. It will be valuable to perform further validation of these results, as more experimental data become available.

Our method enables estimation of log-enrichment in challenging experimental setups comprised of, for example, short reads spanning a sequence of interest; long reads with poor coverage; a mixture of both short and long reads; and settings with more than two conditions—such as when we seek to find sequences enriched for one selection and negatively selected by another, as occurs in engineering gene therapy viral vectors to selectively infect one cell type but not another. In general, our approach also helps to mitigate poor estimates arising from relatively little sequencing data.

Our newly-developed method can immediately leverage any advances in general machine learning classifiers, and naturally handles sequencing reads of variable lengths within a given experiment, so long as the classifier itself does so—as we demonstrated herein using convolutional neural networks. In principle, the predictive performance of such variable-length classifiers can potentially be further improved by incorporating other informative inputs— in addition to read sequence. For example, in settings where it is possible to align to a known reference sequence before modeling, one may supply the mapped position for each read as an additional input. We anticipate that, as high-throughput selection experiments and sequencing-based assays continue to become more varied in their applications, the full potential of model-based enrichment will be further revealed.

## Methods

### Log-enrichment regression

Subjecting two sequence libraries—one for each of two conditions *A* and *B*—to high-throughput sequencing yields a dataset

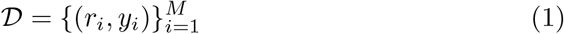

where *r*_*i*_ is the *i*^th^ read’s sequence and *y*_*i*_ is a binary −1*/* + 1 label indicating whether read *r*_*i*_ arose from condition *A* or *B*, respectively. In our analyses of high-throughput selection experiments, the conditions *A* and *B* correspond to pre- and post-selection, but the following methodology applies broadly to settings with sequencing data from two conditions for which we seek to understand or predict sequence properties. In subsequent sections, we also further generalize to more than two conditions.

From these data, 𝒟, one often calculates a count-based log-enrichment (cLE) estimate for each unique sequence [2, 3, 5, 10, 11], which serves as an estimate of the extent to which the sequence has the property being investigated. In selection experiments, we refer to the selection process as acting according to a particular *fitness*, and the cLE estimate thus serves as a proxy for this fitness. To compute cLE estimates, it is convenient to represent 𝒟 in terms of unique sequences: 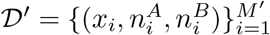 where 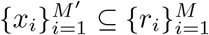 the set of unique observed sequences,

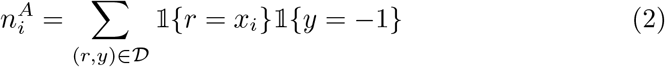

is the observed read count for sequence *x*_*i*_ in the sequencing data for condition *A*, and

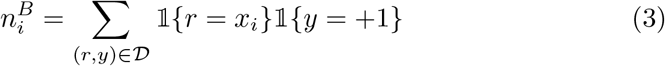

is the corresponding condition *B* read count. For each sequence, the cLE estimate is equal to the log-ratio of read frequencies for conditions *A* and *B*:

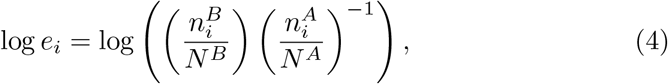

where 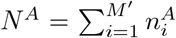 and 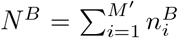. In practice, it is common to add a small constant to each count prior to calculating cLE estimates for mathematical convenience [2, 5]. These “pseudo-counts” stabilize the cLE estimates, and allow one to avoid division by zero for sequences observed in only one condition. In our experiments, we added a pseudo-count of 1 to each raw count.

Log-enrichment (LE) regression approaches fit a model that maps from *x*_*i*_ to log *e*_*i*_. In particular, Zhu et al. [20] derive a weighted least squares procedure for fitting such a regression model; their procedure assigns a weight, 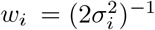, to each sequence, where

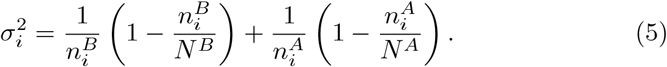

This choice of *w*_*i*_ is motivated by a convergence argument: 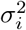 is the asymptotic variance of log *e*_*i*_ [11, 20]. Note that when the counts 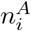 and 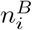 are low, log *e*_*i*_ is a noisier estimate of fitness and the corresponding weight, *w*_*i*_, is smaller. Thus, training a model *f*_*θ*_, with learnable parameters *θ*, using the weighted least squares loss

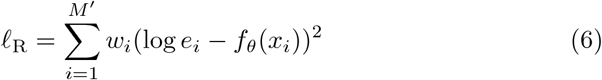

accounts for the heteroscedastic noise in the observed cLE estimates. We refer to this modeling approach as the weighted LE regression (wLER) approach.

### Model-based enrichment

Existing methods that use cLE estimates to train predictive models [20, 21, 29, 32] proceed in two steps: first, one computes a cLE estimate for each observed sequence, and, second, one uses supervised regression to train a model to predict these cLE estimates given the observed sequences. Here, we present a new method, model-based enrichment (MBE) that performs both of these steps at once by reframing LE estimation as a density ratio estimation (DRE) problem. First, we define the density ratio between libraries in each of two conditions and show that a cLE estimate can be viewed as an approximation of the density ratio. Then, we describe the technical details of the MBE approach, which uses a probabilistic classifier trained on sequencing reads to perform DRE.

As in the preceding section, suppose two libraries corresponding to conditions *A* and *B* have been subjected to high-throughput sequencing. Each library can be represented by a discrete probability distribution over sequences: each unique sequence *x*_*i*_ is present in the libraries from conditions *A* and *B* in some ground truth proportions *p*^*A*^(*x*_*i*_), *p*^*B*^(*x*_*i*_) ∈ [0, 1]. The density ratio between these two library distributions is 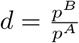.

We can connect this density ratio to cLE estimates. The cLE estimate, log *e*_*i*_, is the log-ratio of the two empirical read frequencies 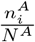 and 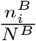 (Eq. 4). These read frequencies are approximations of the true library proportions *p*^*A*^(*x*_*i*_) and *p*^*B*^(*x*_*i*_) based on the observed sample of sequencing reads. Thus, the cLE estimate, log *e*_*i*_, can be viewed as a sample-based approximation of log *d*(*x*_*i*_). LE regression methods can, therefore, be viewed as, first, approximating log *d* using observed counts, and then training a regression model to predict these approximate log-density ratios.

In contrast, DRE techniques [38, 39] can be used to model the density ratio directly from sequencing data. Our proposed MBE approach uses a classification-based DRE technique [36–39] which involves training a probabilistic classifier, *g*_*θ*_, on 𝒟 (Eq. 1) to predict *y*_*i*_ from *r*_*i*_ for each individual read. We use the standard logistic loss,

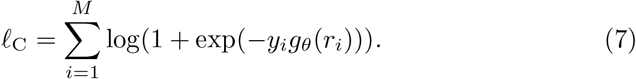

By minimizing this loss with respect to *θ* to obtain the maximum likelihood estimate, 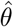, we obtain our predictive model, 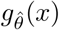 of *p*(*y* | *r* = *x*). By Bayes’ theorem, this also yields an estimator of the density ratio [37, 39]:

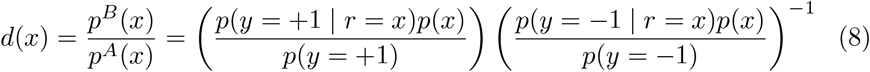

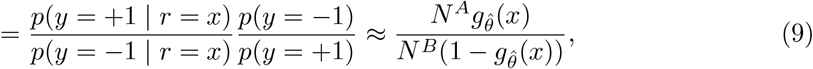

where *N*^*A*^ and *N*^*B*^ are the total read counts for each condition, as in Eq. 4. This ratio of predicted class probabilities, 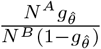, provably converges to *d* [37–39] and is the optimal density ratio estimator among a broad class of semi-parametric estimators (that includes the wLER method) in terms of asymptotic variance under a correctly specified model [37] (Supplementary Note).

MBE naturally accounts for heteroscedastic noise in the observed sequencing data. To see this, we can rewrite *ℓ*_C_ in terms of unique sequences,

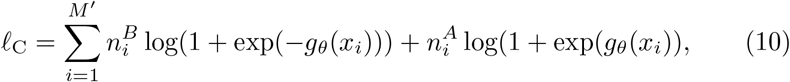

Where 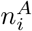 and 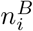 are read counts as defined in Eq. 2-3. This form of *ℓ*_C_ highlights the fact that sequences with higher counts make larger contributions to the loss than those with lower counts, simply by virtue of having been sequenced many times. Thus, *g*_*θ*_ is biased towards modeling *d* more accurately for sequences with more sequencing data, as desired. In this way, the MBE approach accounts for heteroscedasticity in the observed sequencing data without the need to derive a bespoke weighted loss function, unlike the wLER approach.

Note that, in practice, when classes are highly imbalanced (*i. e*., *N*^*A*^ is much larger than *N*^*B*^, or vice versa), the classifier *g*_*θ*_ may implicitly learn to predict the more prevalent class, which may be undesirable. To address this, it may be preferable to use standard machine learning techniques for training classification models under class imbalance, such as class weighting wherein samples from the minority class are up-weighted to have equal overall contribution. Such techniques are not required in our simulations because classes are always exactly balanced (*i. e*., *N*^*A*^ = *N*^*B*^). In our experiments on real sequencing data, we use class weighting (*e. g*., by weighting reads form condition *A* by 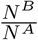 when *A* is the minority class).

### Multi-output modeling

In practice, one often aims to compare sequences across more than two conditions. For example, one may wish to perform multiple rounds of selection for a property of interest (*e. g*., [17]) or to select for multiple different properties (*e. g*., [8]). Here, we describe generalizations of the MBE and wLER approaches that can be used to model high-throughput sequencing data collected from more than two conditions. In this setting, one has sequencing data 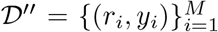 where *r*_*i*_ is the *i*^th^ read’s sequence and *y*_*i*_ is a categorical label indicating the condition from which the read *r*_*i*_ arose. For example, if one runs an experiment selecting for *k* ∈ **N** different properties, one can define *y* ∈ {0, 1, …, *k*} where *y*_*i*_ = 0 indicates a read from the pre-selection sequencing data, and *y*_*i*_ = *j* for *j* ∈ { 1, …, *k*} indicates a read from the post-selection sequencing data for the *j*^th^ property.

It is straightforward to handle multiple conditions using the MBE approach: instead of using a binary classifier, one trains a multi-class classification model, *g*_*θ*_, to predict the categorical label *y*_*i*_ from read sequence *r*_*i*_ using a standard categorical cross-entropy loss. This produces a model of *p*(*y* | *r*) which can be used to estimate the density ratios 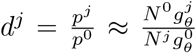, where *p*^*j*^ denotes the true probability distribution corresponding to the library in the *j*^th^ condition and 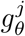 denotes the predicted class probability for *y* = *j*.

For the wLER approach, the data 𝒟′ can be converted into cLE estimates for each unique sequence:

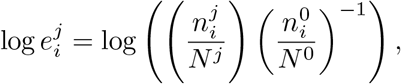

where 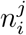 is the number of times the sequence *x*_*i*_ appeared in the sequencing data for the *j*^th^ condition and *N*^*j*^ is the total number of reads from the *j*^th^ condition. One can, then, fit a multi-output regression model that jointly predicts the cLE estimates for each condition from sequence. The overall loss for training a such a multi-output model, *f*_*θ*_, using wLER is

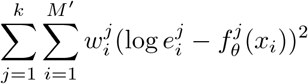

where 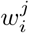 is the weight for the *i*^th^ sequence and *j*^th^ condition, and 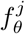 denotes the *j*^th^ model output.

### Model architectures and training

In our experiments, we aim to compare and contrast the general performance of the wLER and MBE approaches across a broad range of settings. To enable direct comparison of the two methods, we implemented wLER and MBE using the same model architectures and hyperparameters for the underlying regression and classification models. We will, next, describe each model architecture and provide implementation details.

We used eleven different model architectures: seven architectures that are the same as in Zhu *et al*. [20]—three linear models and four fully-connected neural networks (NNs)—and four convolutional neural network (CNN) models that can operate on variable-length inputs. The linear models each use one of three input representations: (1) an “independent site” (IS) representation comprised of one-hot encodings of individual amino acids, (2) a “neighbor” representation comprised of the IS features and one-hot encodings of the pairwise interactions between pairs of positions directly adjacent in sequence, and (3) a “pairwise” representation comprised of the IS features and one-hot encodings of all possible pairwise interactions. All NNs use IS input features and have two hidden layers, and differ by the number of hidden units: 100, 200, 500, or 1000 units per layer. The CNNs differ in the number of convolutional layers used (2, 4, 8, or 16), but all use IS input features, convolutions with a window of size 5 and 100 filters, residual and skip connections, and a global max pooling layer as the penultimate layer.

All models were trained using the AMSGrad Adam optimizer [49] with default learning rate (10^−3^) for ten epochs. For the linear models and NNs, we used the default value for Adam’s *ϵ* parameter (10^−7^); for the CNNs, we set *ϵ* = 10^−4^ and applied gradient clipping with a threshold of 1. We performed three-fold cross-validation at the sequence level: for each fold, one third of the unique sequences in the library (and their corresponding sequencing reads) were held-out as a test set.

### Simulating ground truth fitness

We constructed several simulated datasets to help analyze the strengths and weaknesses of MBE, wLER, and cLE across different practical settings. These simulations were motivated by high-throughput selection experiments [8, 20, 31] which perform a selection on large sequence libraries for a property of interest, such as fluorescence [31]. To simulate such selection experiments, we first simulate the ground truth *fitness function* that maps sequence to property, then use this fitness to simulate selection. In the remainder of this section, we describe the process used to simulated fitness as a linear function of independent amino acid sites and randomly selected higher-order epistatic interactions. In the following section, we describe the procedure to simulate selection using simulated fitness.

First, we give a brief overview of the process used to simulate ground truth fitness before providing the technical details. For a given sequence of interest, we first constructed a set containing all independent amino acid sites and a user-specified number of combinations of sites—such as an epistatic combination of the second, third, and tenth positions—drawn randomly from among all possible higher-order epistatic interactions between positions. The degree of each epistatic effect (2 up to the sequence length) is drawn randomly based on an empirical estimate of this degree distribution. The fitness function is, then, taken to be a linear function of all the independent sites and epistatic terms in this constructed set with random coefficients.

In more detail, for a sequence *x* of length *L*_*a*_ amino acids, we simulated the fitness function, *F*_*T*_ (*x*) as

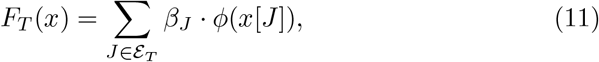

where *T* is the hyper-parameter controlling the maximum number of epistatic terms included in *F*_*T*_ ;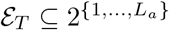, is a set of index sets—each of which represents an independent site or a particular higher-order epistatic combination—whose construction is described below; *x*[*J*] is the subsequence of *x* at the positions in the index set *J*; *ϕ* denotes standard one-hot encoding; and the coefficients are sampled according to *β*_*J*_ ∼ 𝒩 (**0**, 2^−|*J*|^**I**).

We constructed *ε*_*T*_ (the specific set of first-order and higher-order epistatic terms to include in the simulated fitness function) to contain all singleton sets ({{*i*} | *i* ∈ {1, …, *L*_*a*_}} ⊆ *ε*_*T*_), so that *F*_*T*_ includes terms for all independent sites. In addition, *ε*_*T*_ contains *T* randomly-chosen non-singleton index sets, each generated by:

1. randomly choosing the order of epistasis, *R*, by sampling 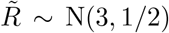 (based on visual inspection of the empirical bell-shaped distribution of the orders of statistically significant epistatic terms in Poelwijk *et al*.. [29]), and taking 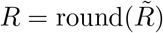; and
2. choosing the specific positions included in the epistatic term by sampling *R* times without replacement from {1, …, *L*_*a*_}.

To guide our choice of *T*, we combined the following insights: (i) for a fluorescent protein with 13 amino acids, 260 epistatic terms are sufficient for an accurate model of fitness [29]; (ii) the number of contacts in a protein scales linearly with sequence length [43, 44]; and (iii) recent work suggests that the sparsity of higher-order epistatic interactions in fitness landscapes is closely related to structural contact information [45]. We, therefore, hypothesized that the linear scaling 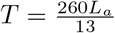 provides a reasonable starting point for analyses.

### Simulating preand post-selection sequencing data

The wLER and MBE approaches both aim to accurately quantify sequences of interest based on high-throughput sequencing data. We used simulated highthroughput selection datasets to compare each method’s ability to quantify sequences accurately using sequencing data, which requires simulating sequencing reads from pre- and post-selection libraries. Here, we detail the process of simulating sequencing reads given library sequences and a ground truth fitness function. Then, in the subsequent sections, we will describe how we combined this process with three specific approaches for simulating library sequences to construct our datasets.

Let 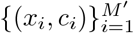 be pairs of, respectively, a unique library sequence and its true count—as generated, for example, by one of the three library construction simulations described in the subsequent sections. In addition, let *F*_*T*_ be a ground truth fitness function simulated as in the previous section. Briefly, the process to simulate sequencing reads from pre- and post-selection libraries proceeds as follows: first, we generate a pre-selection library distribution by adding a small random perturbation to the empirical distribution 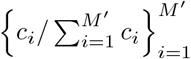. This step simulates slight distributional perturbations that may occur with PCR amplification, and also has the nice side-effect of allowing one to generate multiple replicates with slightly different pre- and post-selection library distributions for the same set of unique sequences 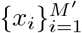. Next, we simulate selection according to the fitness *F*_*T*_ : the post-selection library distribution is determined by scaling the pre-selection distribution using 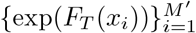, which ensures that the ground truth log-density ratio is proportional to the specified fitness 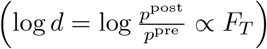. Finally, we sample from the pre- and post-selection distributions to simulate sequencing reads, optionally truncating each read to 100 amino acids uniformly at random to generate short reads.

In more detail, we simulated pre- and post-selection sequencing data by:

1. sampling 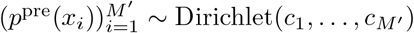;
2. setting

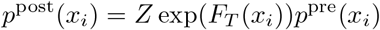

where 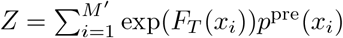 is a normalization constant;
3. sampling pre- and post-selection sequencing counts according to

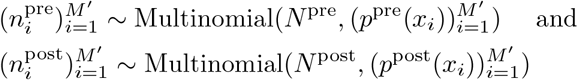

for some desired number of sequencing reads, *N*^pre^ and *N*^post^; and, if simulating short reads, additionally
4. sampling 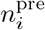 and 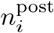 contiguous 100-mers from *x*_*i*_ uniformly at random.

### Simulated insertion libraries

To empirically compare and contrast our MBE approach to the wLER approach in practical settings, we sought to simulate realistic sequence libraries motivated by experimental constructions from recent studies.

We simulated diversified libraries of insertion sequences motivated by our work in adeno-associated virus (AAV) capsid engineering [20]. In this study, we used a library of 21-mer nucleotide insertion sequences, where each codon was independently sampled from the distribution defined by the NNK degenerate codon: “NN” denotes a uniform distribution over all four nucleotides in the first two positions of a codon and “K” denotes equal probability on nucleotides G and T in the third codon position. Here, we sampled sequences from this NNK distribution to simulate three insertion libraries containing length 21, 150, and 300 nucleotide sequences, respectively. Specifically, each sequence is generated by sampling either 7, 50, or 100 codons independently from the NNK distribution. To keep each of our simulated insertion datasets as similar as possible to the experimental data from Zhu *et al*. [20], we sampled sequences in this manner until we obtained a set of 8.5 × 10^6^ unique library sequences. We take the set 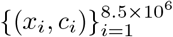 to be the simulated library, where *x*_i_ is the *i*^th^ unique insertion sequence and *c*_*i*_ is the number of times it was sampled from the NNK distribution before 8.5 × 10^6^ unique sequences were generated.We used *T* = 140, 1000, and 2000 to simulate ground truth epistatic fitness for the 21-mer, 150-mer, and 300-mer insertion libraries, respectively, and simulated *N*^pre^ = *N*^post^ = 4.6 × 10^7^ sequencing reads for each library using the process described in the previous section.

To gain insight into the effect of sequencing error on MBE and wLER, we also constructed a noisy version of the sequencing data for the 21-mer insertion library containing simulated sequencing errors in both the pre- and post-selection sequencing reads. Because Illumina’s next-generation sequencers have an approximately 1% error rate and predominantly produce substitution errors [47], we added substitution errors to each position of each simulated read uniformly at random with probability 0.01.

### Simulated avGFP mutagenesis library

Motivated by a recent study of the fitness landscape of the green fluorescent protein from *Aequorea victoria* [31], we generated an avGFP library by mutating positions of the avGFP reference sequence from Sarkisyan et al. [31] (238 amino acids long) uniformly at random. We used a mutation rate of 10% to generate 2.5 × 10^7^ unique library sequences. Specifically, we generated mutated avGFP sequences—by mutating each position independently with probability 0.01—until we obtained a set 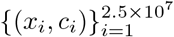, where each *x*_i_ a unique library sequence and *c*_*i*_ is the number of times it was generated before 2.5 × 10^7^ unique sequences were obtained.

To simulate selection and sequencing, we used *T* = 4, 760 to simulate ground truth fitness, and generated both long-read (*N*^pre^ = *N*^post^ = 4.6 × 10^5^ to be within PacBio’s throughput [34, 35]) and short-read (*N*^pre^ = *N*^post^ = 4.6 × 10^7^ to match the dataset from Zhu *et al*. [20]) sequencing data.

### Simulated AAV recombination library

We simulated a recombination library of AAV capsid sequences motivated by an AAV directed evolution study [8], wherein several AAV serotypes are recombined using seven crossovers separating eight recombination blocks. We generated library sequences by recombining AAV serotypes 1-9 with seven uniformly-spaced crossovers. This library contains 26,873,856 unique library sequences that are 2,253 nucleotides long. We simulated epistatic fitness with *T* = 15, 020.

To assess the effects of the type and amount of sequencing data, we generated multiple datasets: three long-read datasets with *N*^pre^ = *N*^post^ = 4.6 × 10^3^, 4.6 × 10^4^, and 4.6 × 10^5^, respectively; one short-read dataset with *N*^pre^ = *N*^post^ = 4.6 × 10^7^; and one hybrid dataset containing 4.6 × 10^3^ long reads and 4.5 × 10^7^ short reads for both pre- and post-selection. To help gain insights into the effects of sequencing error, we also constructed a noisy AAV recombination dataset that incorporated simulated sequencing errors into 4.6 × 10^5^ pre- and post-selection sequencing reads using SimLoRD [48] to simulate PacBio SMRT sequencing errors.

## Declarations

## Availability of data and materials

Code for generating simulated datasets, training models, and evaluating performance will be made available upon publication.

## Funding

AB’s work was supported by the National Science Foundation Graduate Research Fellowship Program under Grant No. 1752814. Any opinions, findings, and conclusions or recommendations expressed in this material are those of the author(s) and do not necessarily reflect the views of the National Science Foundation. JL’s work was supported in part by a Chan Zuckerberg Investigator award.

## Supplementary Figures

**Supplementary Fig. 1.**
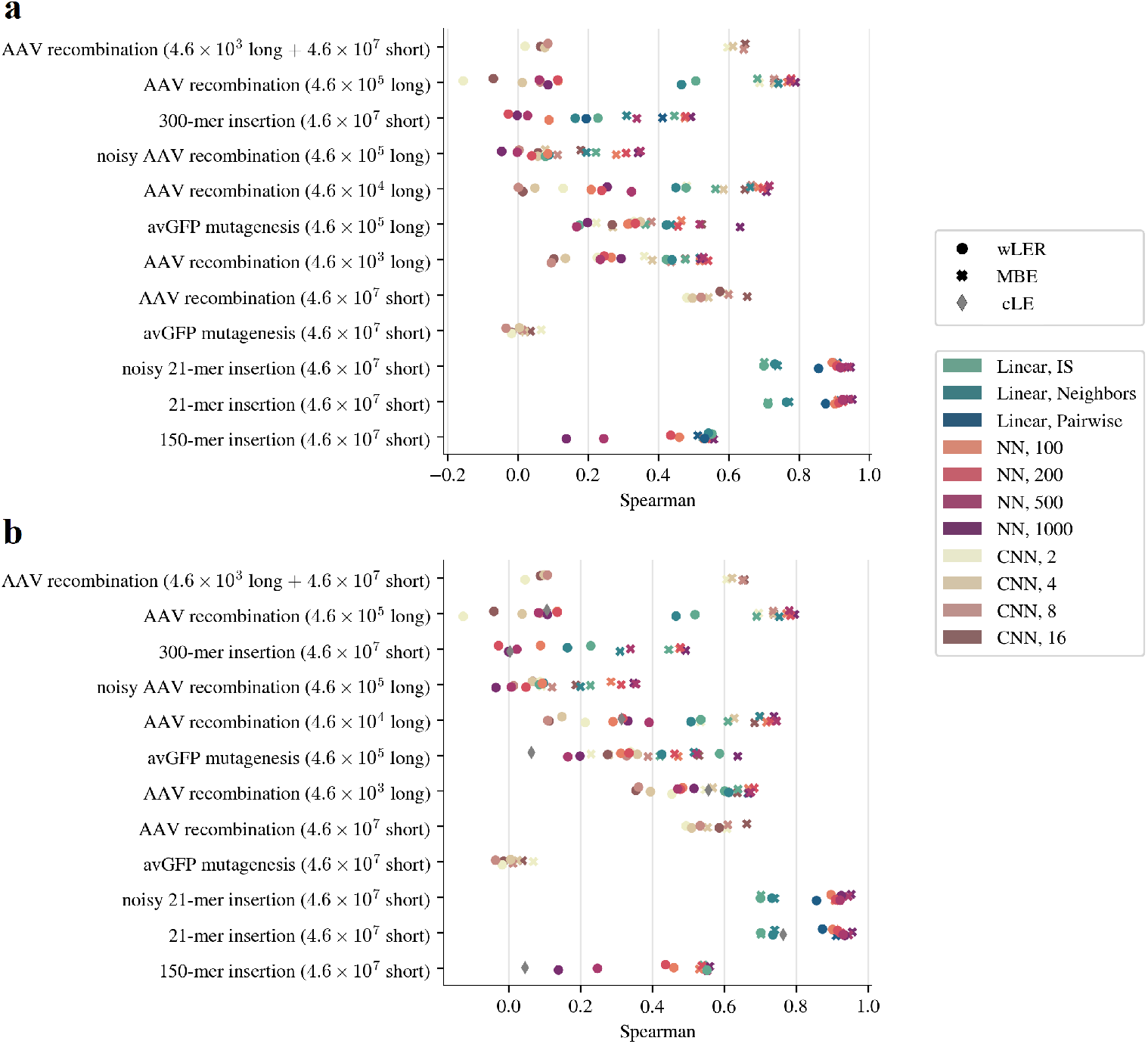
Simulation results for all model architectures. (a) and (b) are the same as Fig. 1a and b, respectively, but display the Spearman correlation between model predictions and ground truth fitness for all model architectures.

**Supplementary Fig. 2.**
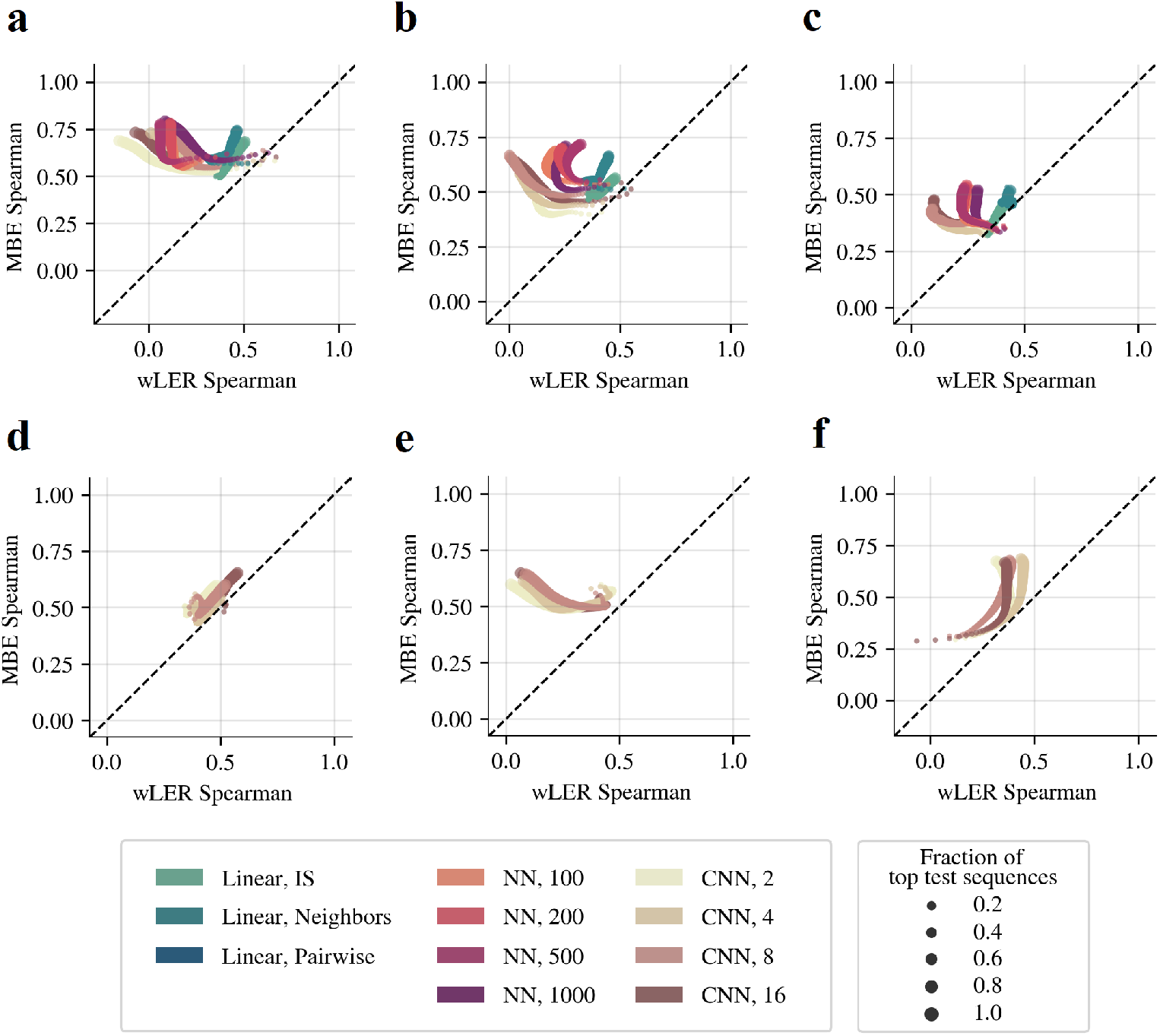
Simulated library results with increasing long read sparsity and short reads. Compares Spearman correlation between simulated ground truth fitness and wLER or MBE predictions on held-out sequences of interest when models are trained using the simulated AAV recombination datasets with (a) 4.6 × 10^5^ long reads, (b) 4.6 × 10^4^ long reads, (c) 4.6 × 10^3^ long reads, (d) 4.6 10^7^ short reads, and (e) a combination of 4.6 × 10^3^ long and 4.6 × 10^7^ short reads, and (f) the avGFP mutagenesis dataset with 4.6 × 10^7^ short reads. Each panel compares the Spearman correlation achieved by the wLER and MBE approaches using the same model architecture and hyper-parameters. Dot size represents the fraction of test sequences with highest ground truth fitness used to compute Spearman correlation. Only CNNs are included in d-f since the linear and NN models cannot operate on variable-length sequences.

**Supplementary Fig. 3.**
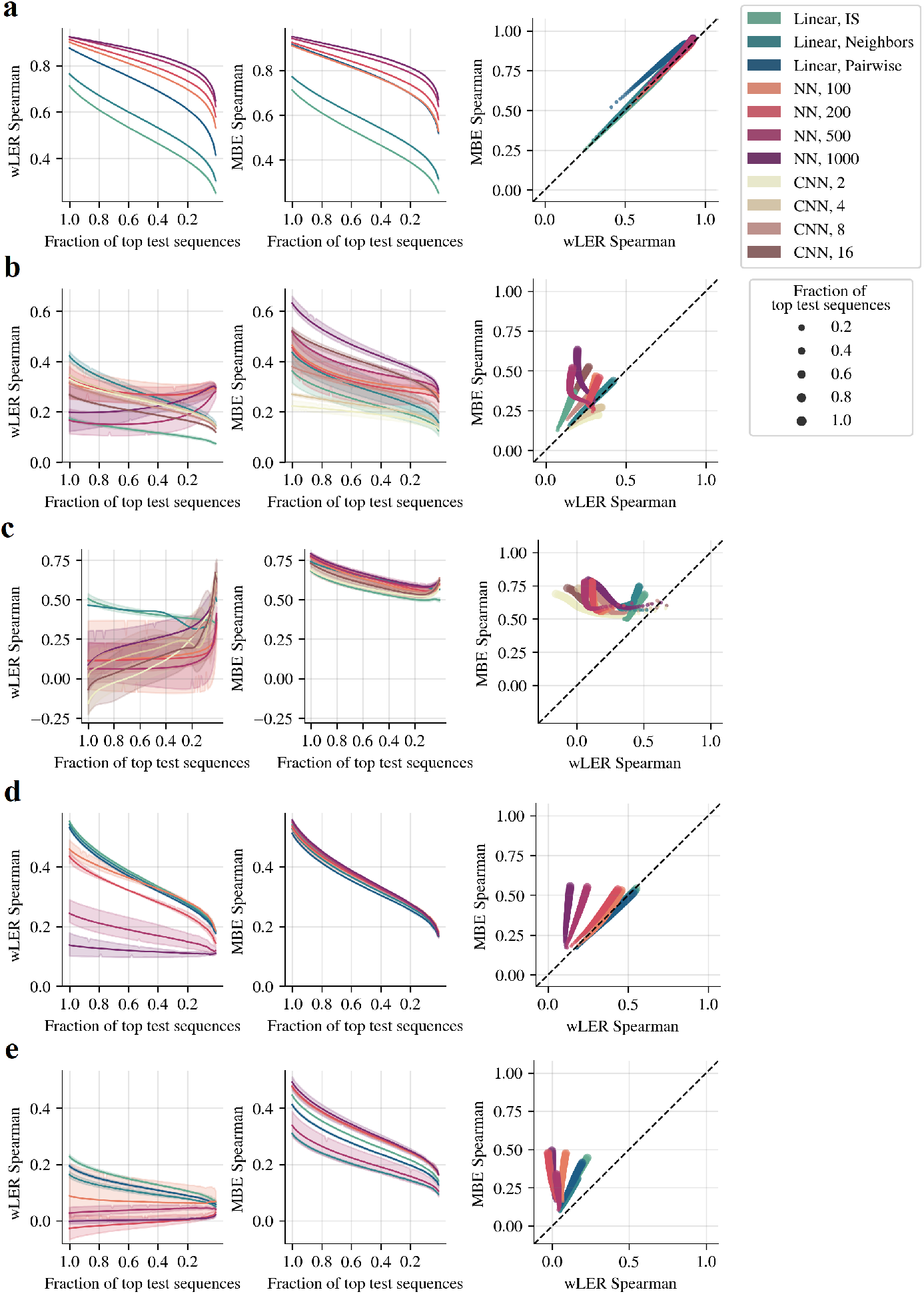
Generalized Spearman for simulated library prediction. Comparison of generalized Spearman correlation between simulated ground truth fitness and wLER or MBE predictions on held-out full-length sequences for the simulated (a) 21-mer insertion (4.6 × 10^7^ short reads), (b) avGFP mutagenesis (4.6 × 10^5^ long reads), (c) AAV recombination (4.6 × 10^5^ long reads), (d) 150-mer insertion (4.6 × 10^7^ short reads), and (e) 300-mer insertion (4.6 × 10^7^ short reads) datasets. In each row, the leftmost panel compares the performance of wLER for each model architecture (the horizontal axis displays the fraction of top test sequences with highest ground truth fitness used to calculate Spearman correlation), the center panel is the same as the left panel MBE, and the rightmost panel is a paired plot version of the left and center panels (dot size represents the fraction of top test sequences used to compute Spearman correlation).

**Supplementary Fig. 4.**
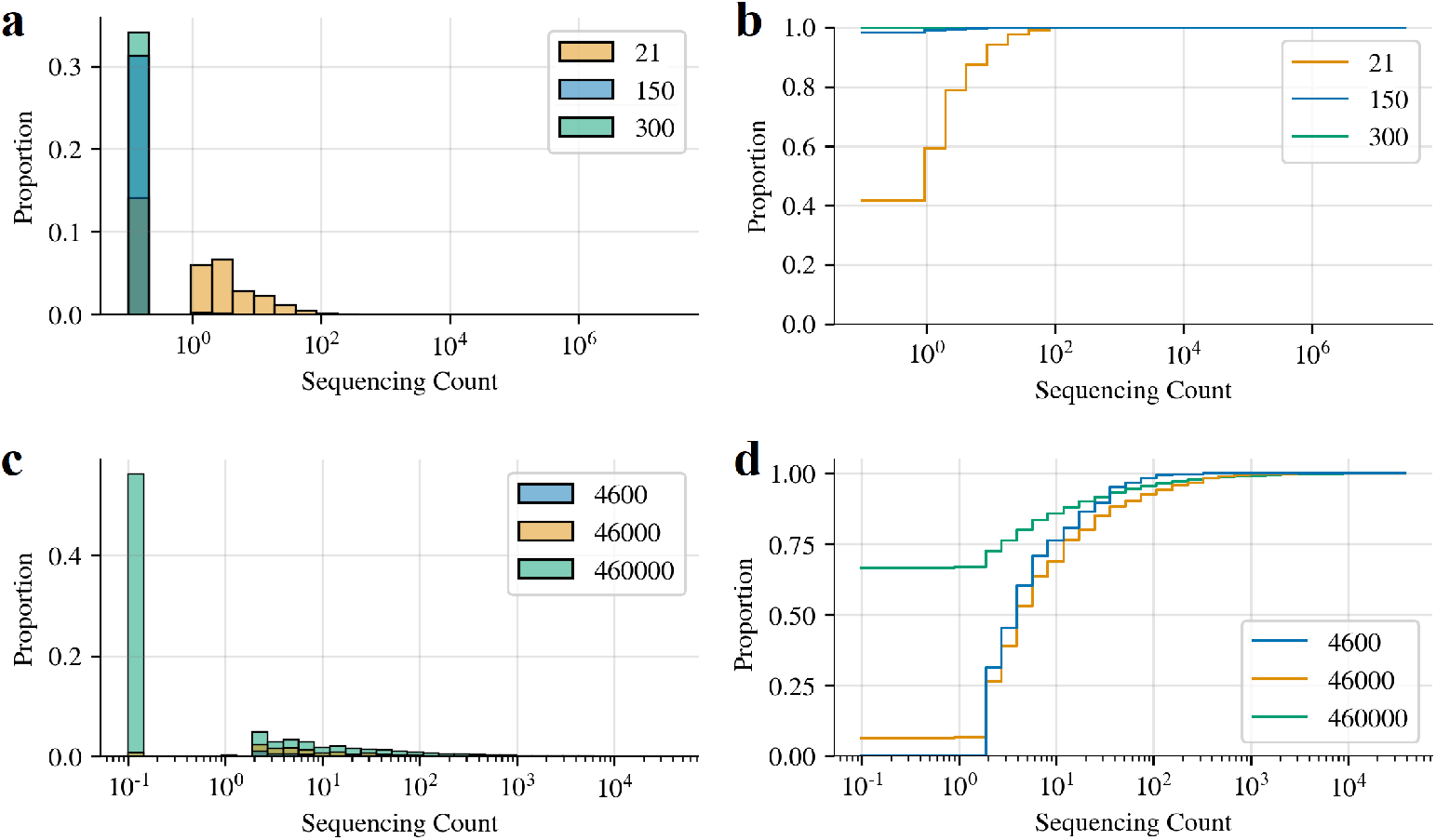
Sequencing count histograms for simulated insertion and recombination libraries. Histogram (left) and cumulative histogram (right) of simulated post-selection sequencing counts for the (a-b) 21-mer, 150-mer, and 300-mer insertion datasets, and (c-d) AAV recombination datasets with 4.6 × 10^5^, 4.6 × 10^4^, and 4.6 × 10^3^ long reads.

**Supplementary Fig. 5.**
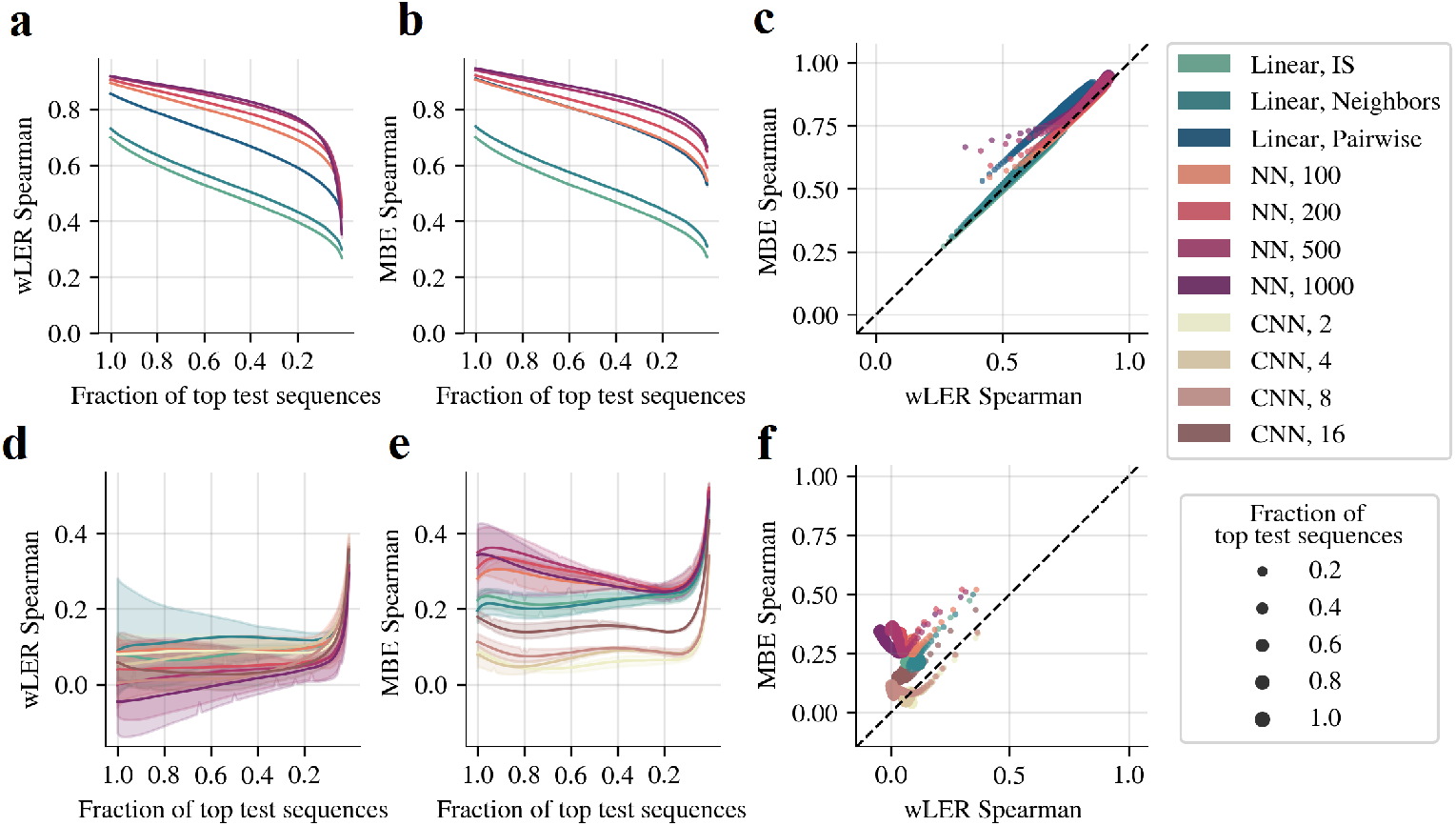
Generalized Spearman for prediction with simulated sequencing errors. Comparison of the Spearman correlation between simulated ground truth fitness and wLER or MBE predictions on held-out full-length library sequences when models are trained using the simulated (a-c) noisy 21-mer insertion (4.6 × 10^7^ short reads) and (d-f) noisy AAV recombination (4.6 × 10^5^ long) datasets. The noisy 21-mer insertion dataset includes substitution errors added to the training set at a uniform error rate of 1%, consistent with Illumina’s next-generation sequencers [47]. The noisy AAV recombination dataset contains simulated PacBio SMRT sequencing errors added to the training set using SimLoRD [48]. In each row, the leftmost panel compares the performance of wLER for each model architecture (the horizontal axis displays what fraction of top test sequences with highest ground truth fitness is used to calculate Spearman correlation), the center panel is the same as the left panel for MBE, and the rightmost panel is a paired plot version of the left and center plots (dot size represents the fraction of top test sequences used to compute Spearman correlation).

**Supplementary Fig. 6.**
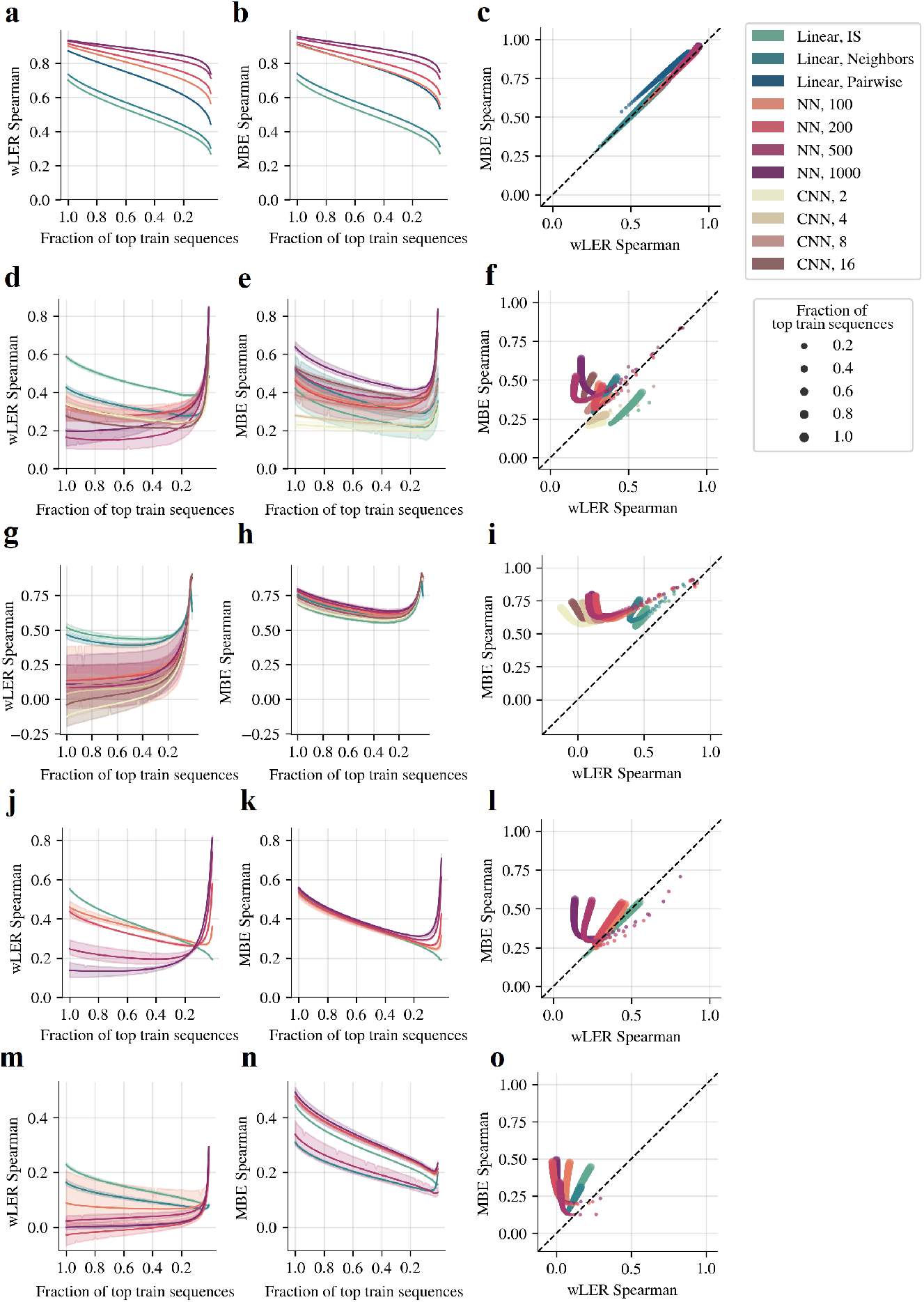
Generalized Spearman for simulated library estimation. Comparison of the Spearman correlation between simulated ground truth fitness and wLER or MBE LE estimates for full-length library sequences observed during training for the simulated (a-c) 21-mer insertion (4.6 × 10^7^ short reads), (d-f) avGFP mutagenesis (4.6 × 10^5^ long reads), (g-i) AAV recombination (4.6 × 10^5^ long reads), (j-l) 150-mer insertion (4.6 × 10^7^ short reads), and (m-o) 300-mer insertion (4.6 × 10^7^ short reads) datasets. In each row, the leftmost panel compares the performance of wLER for each model architecture (the horizontal axis displays what fraction of top test sequences with highest ground truth fitness is used to calculate Spearman correlation), the center panel is the same as the leftmost panel for MBE, and the rightmost panel is a paired plot version of the left and center plots (dot size represents the fraction of top test sequences used to compute Spearman correlation).

**Supplementary Fig. 7.**
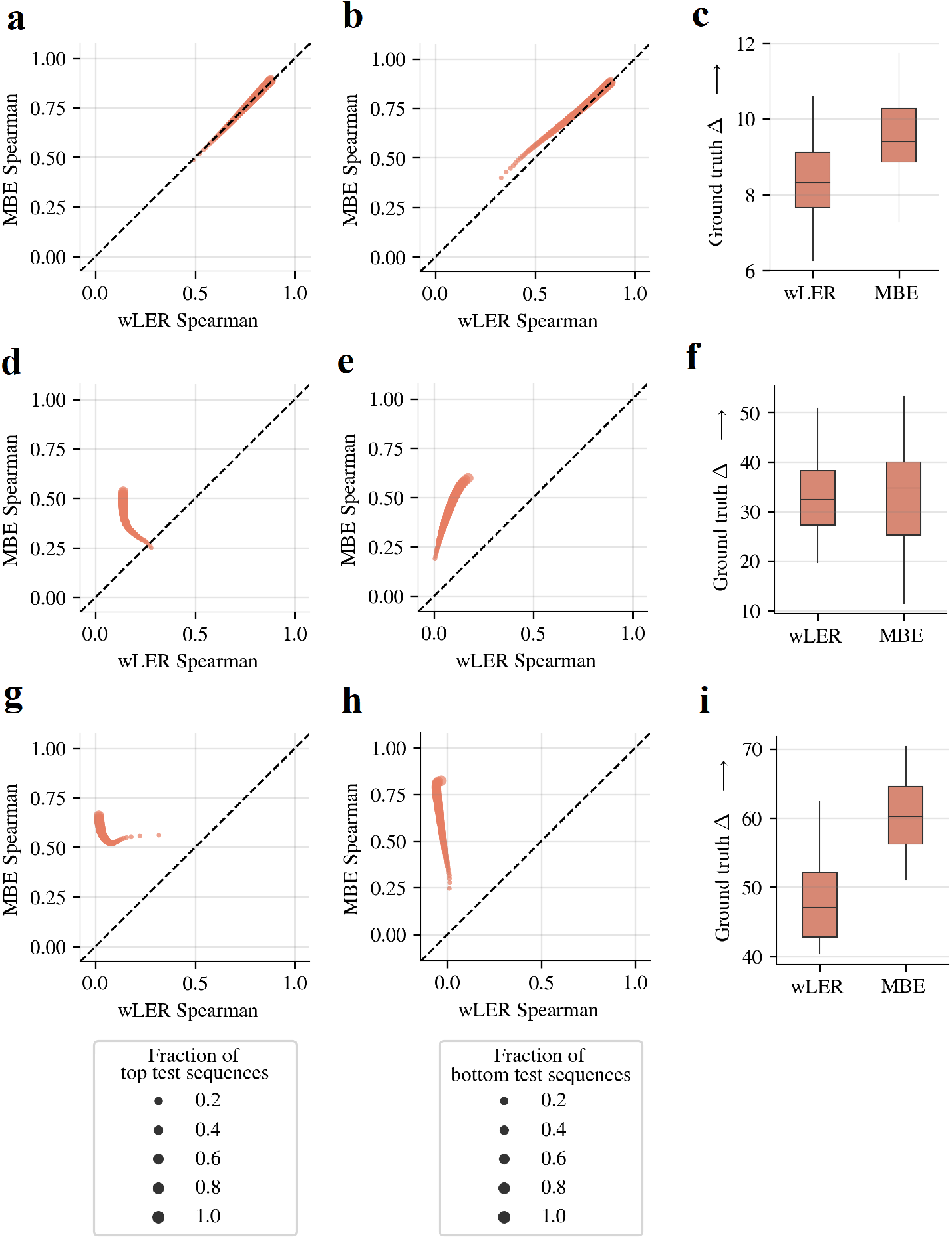
Simulated positive, negative, and selectivity selection results. Comparison of wLER and MBE on (left) prediction for sequences with high ground truth positive fitness, (center) prediction for sequences with low ground truth negative fitness, and (right) selection for sequence selectivity for the simulated (a-c) 21-mer insertion (4.6 × 10^7^ short reads), (d-f) avGFP mutagenesis (4.6 × 10^5^ long reads), and (g-i) AAV recombination (4.6 × 10^5^ long reads) datasets. For positive fitness, dot size represents the fraction of top test sequences according to highest ground truth positive fitness. For negative fitness, dot size represents the fraction of test sequences with lowest ground truth negative fitness. In each row, the rightmost panel displays ground truth selectivity (the difference between positive and negative fitness values, Δ) for the top ten test sequences according to each model’s predicted selectivity (the difference between predicted fitness values) for each of the three test folds.

**Supplementary Fig. 8.**
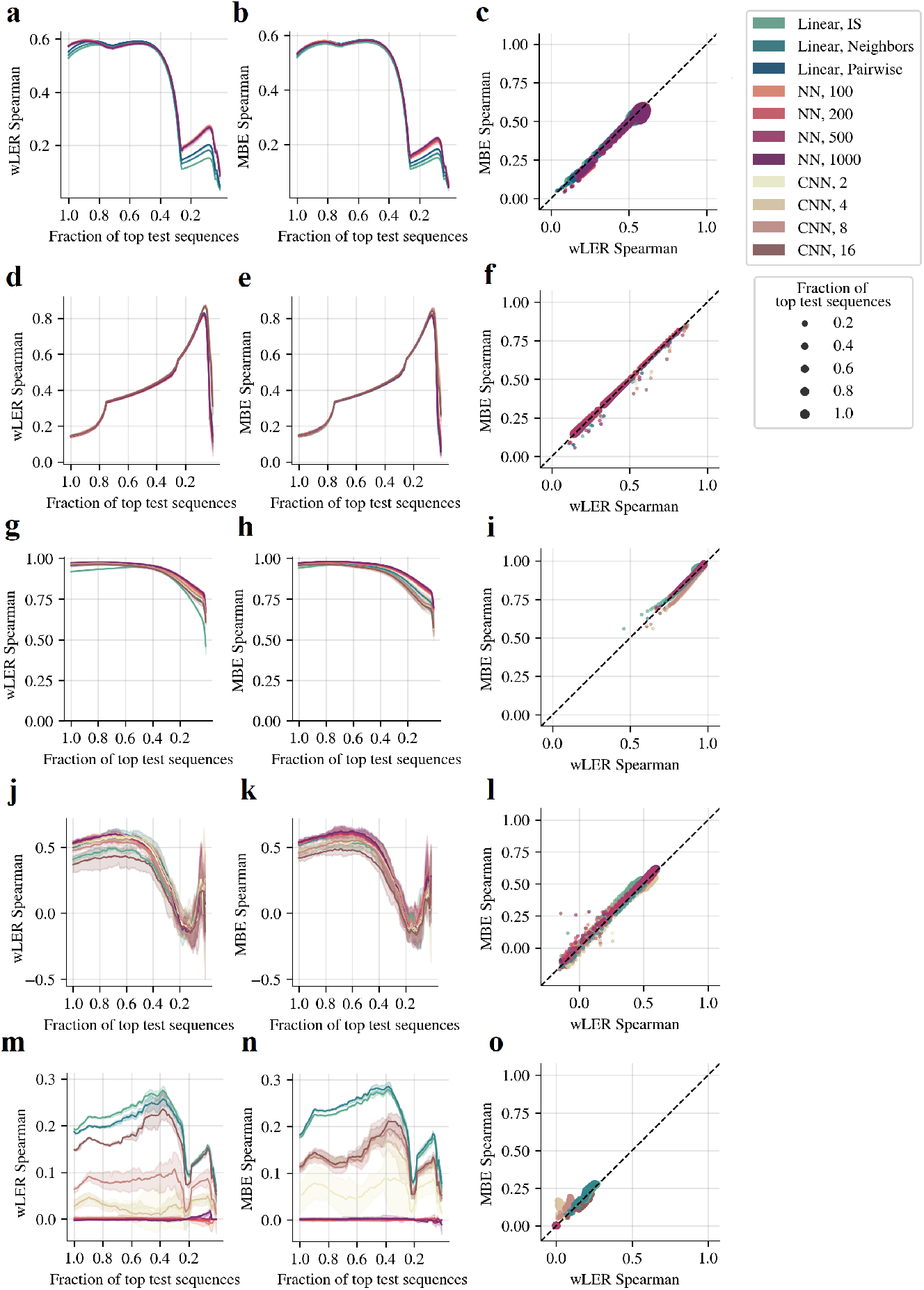
Experimental library cross-validation results. Comparison of the Spearman correlation between wLER or MBE predictions and observed cLE estimates on the real sequencing datasets from (a-c) the AAV5 insertion library from Zhu *et al*. [20], (d-f) the SARS-CoV-2 tiled peptide library from Huisman *et al*. [17], (g-i) the GB1 double site saturation mutagenesis library from Olson *et al*. [12], (j-l) the library of natural and designed chorismate mutase homologs from Russ *et al*. [28], and (m-o) the Bgl3 random mutagenesis library from Romero *et al*. [27]. In each row, the leftmost panel compares the performance of wLER for each model architecture restricted to a given top fraction of heldout test sequences with highest observed cLE estimate, the center panel is the same as the leftmost panel for MBE, and the rightmost panel is a paired plot version of the left and center panels (dot size represents the fraction of top test sequences with highest observed cLE estimate used to compute Spearman correlation).

**Supplementary Fig. 9.**
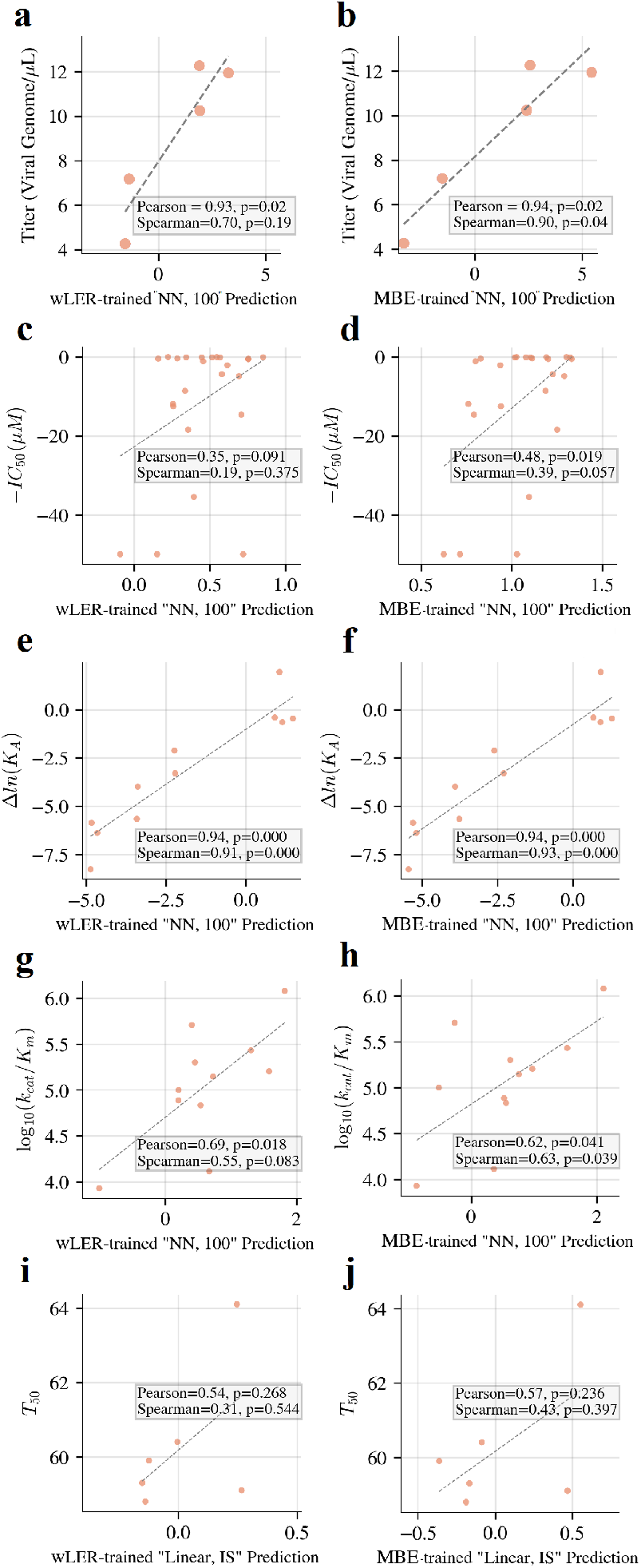
Low-throughput experimental property measurement predictions. Comparison of (left) wLER and (right) MBE predictions and experimental property measurements from (a-b) Zhu *et al*. [20] (packaging titer), (c-d) Huisman *et al*. [17] (*IC*_50_, half maximal inhibitory concentration), (e-f) Olson *et al*. [12] (Δln(*K*_*A*_), change in log-binding constant), (g-h) Russ *et al*. [28] (log_10_(*k*_*cat*_*/K*_*m*_), log-second-order reaction rate constant), and (i-j) Romero *et al*. [27] (*T*_50_, temperature where half of the protein is inactivated in ten minutes). The 100-unit NN architecture is used for all datasets except that from Romero *et al*. [27] for which the linear architecture with IS features is used.

**Supplementary Table 1.**
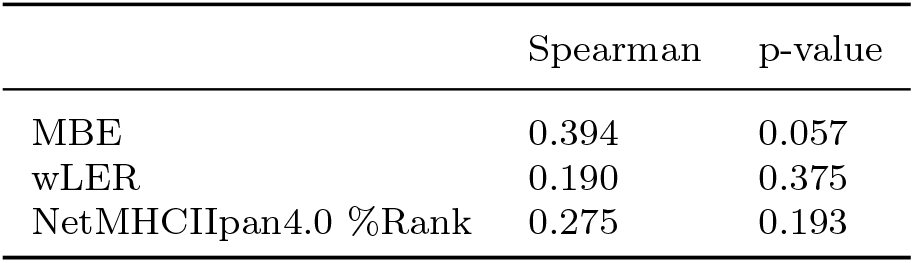
Comparison of Spearman correlation between experimental *IC*_50_ measurements from Huisman *et al*. [17] and wLER predictions, MBE predictions, or reported NetMHCIIpan4.0 predictions from Huisman *et al*. [17]. The 100-unit NN architecture is used for the wLER and MBE methods.

## Supplementary Note

### Asymptotic optimality of model-based enrichment

In this section, we review key parametric convergence results which imply that, under the assumption of a correctly specified parametric model, the proposed model-based enrichment (MBE) estimator is optimal among a broad class of semi-parametric density ratio estimators—including the weighted log-enrichment regression (wLER) method [20]—in terms of asymptotic variance. We begin by recalling some notation: let *p*^*A*^ and *p*^*B*^ be two probability distributions, 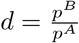 be their density ratio, and

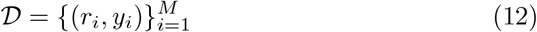

be a dataset of observed samples where *y*_*i*_ is a binary label indicating whether the sample *r*_*i*_ is from *p*^*A*^ (*y*_*i*_ = − 1) or *p*^*B*^ (*y*_*i*_ = +1). Further, let *N*^*A*^ and *N*^*B*^ be the number of samples from *p*^*A*^ and *p*^*B*^, respectively. Recall that the MBE approach uses logistic regression to learn a classifier that predicts *p*(*y*_*i*_ | *r*_*i*_), and these predicted class probabilities give an estimate of the density ratio (Methods). In other words, the MBE approach estimates the density ratio using the parametric model

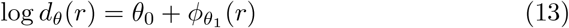

where *θ*_0_ ∈ **R**, *θ* = (*θ*_0_, *θ*_1_) ∈ **R**^*b*^ is a *b*-dimensional parameter, and *ϕ*_*θ*1_ real-valued function (*e. g*., defined by the choice of model architecture).

Whenever correctly-specified density models for both *p*^*A*^ and *p*^*B*^ are unavailable, direct density ratio estimation of 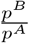 —as performed by the MBE approach—is preferable compared to separate density estimation of *p*^*A*^ and *p*^*B*^ in terms of asymptotic unnormalized Kullback–Leibler divergence to the true density ratio, *d* [39]. Moreover, Qin [37] showed that, if the logistic regression model is correctly specified—that is, if the true density ratio *d* is realized by *d*_*θ*_^*^ in the parametric model—then the MBE approach is optimal among a large class of semi-parametric density ratio estimators in the sense that it has the smallest asymptotic variance. Specifically, the class of semi-parametric estimators in Qin’s analysis is a class of generalized moment-matching estimators:

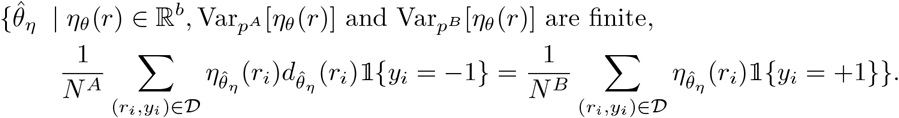

This class of estimators contains several popular density ratio estimators, including the Kullback-Leibler (KL) importance estimation procedure [38, 39] that learns a density ratio model by minimizing empirical KL divergence between *d* · *p*^*A*^ and *p*^*B*^. Other estimation techniques, including weighted and non-linear least squares regression, can also be cast in terms of generalized moment-matching optimization [50] and, therefore, the wLER approach is included in Qin’s class of estimators, as are several other existing log-enrichment regression approaches [19, 29]. Thus, under a correctly specified parametric model, the MBE approach is the preferred density ratio estimation technique—and, in the context of this work, the preferred technique for quantifying sequences based on sequencing data from a high-throughput screen or selection—in terms of asymptotic variance.

## Notes

### Competing Interest Statement

The authors have declared no competing interest.

### Summary of Updates

Section on Model-based enrichment updated to clarify derivation; Figure 3 revised

## References

[1] Lane, M., Seelig, B.: Directed evolution of novel proteins. Curr Opin Chem Biol 22, 129–126 (2014)

[2] Matuszewski, S., Hildebrandt, M.E., Ghenu, A.-H., Jensen, J.D., Bank, C.: A statistical guide to the design of deep mutational scanning experiments. Genetics 204(1), 77–87 (2016)

[3] Wrenbeck, E.E., Faber, M.S., Whitehead, T.A.: Deep sequencing methods for protein engineering and design. Current opinion in structural biology 45, 36–44 (2017)

[4] Trapnell, C., Roberts, A., Goff, L., Pertea, G., Kim, D., Kelley, D.R., Pimentel, H., Salzberg, S.L., Rinn, J.L., Pachter, L.: Differential gene and transcript expression analysis of rna-seq experiments with tophat and cufflinks. Nature protocols 7(3), 562–578 (2012)

[5] Rubin, A.F., Gelman, H., Lucas, N., Bajjalieh, S.M., Papenfuss, A.T., Speed, T.P., Fowler, D.M.: A statistical framework for analyzing deep mutational scanning data. Genome biology 18(1), 1–15 (2017)

[6] Wenger, A.M., Peluso, P., Rowell, W.J., Chang, P.-C., Hall, R.J., Concepcion, G.T., Ebler, J., Fungtammasan, A., Kolesnikov, A., Olson, N.D., et al.: Accurate circular consensus long-read sequencing improves variant detection and assembly of a human genome. Nature biotechnology 37(10), 1155–1162 (2019)

[7] Ogden, P.J., Kelsic, E.D., Sinai, S., Church, G.M.: Comprehensive aav capsid fitness landscape reveals a viral gene and enables machine-guided design. Science 366(6469), 1139–1143 (2019)

[8] Ojala, D.S., Sun, S., Santiago-Ortiz, J.L., Shapiro, M.G., Romero, P.A., Schaffer, D.V.: In vivo selection of a computationally designed schema aav library yields a novel variant for infection of adult neural stem cells in the svz. Molecular Therapy 26(1), 304–319 (2018)

[9] Patwardhan, R.P., Lee, C., Litvin, O., Young, D.L., Pe’er, D., Shendure, J.: High-resolution analysis of dna regulatory elements by synthetic saturation mutagenesis. Nature biotechnology 27(12), 1173–1175 (2009)

[10] Fowler, D.M., Araya, C.L., Gerard, W., Fields, S.: Enrich: software for analysis of protein function by enrichment and depletion of variants. Bioinformatics 27(24), 3430–3431 (2011)

[11] Katz, D., Baptista, J., Azen, S., Pike, M.: Obtaining confidence intervals for the risk ratio in cohort studies. Biometrics, 469–474 (1978)

[12] Olson, C.A., Wu, N.C., Sun, R.: A comprehensive biophysical description of pairwise epistasis throughout an entire protein domain. Current biology 24(22), 2643–2651 (2014)

[13] Robinson, M.D., McCarthy, D.J., Smyth, G.K.: edger: a bioconductor package for differential expression analysis of digital gene expression data. bioinformatics 26(1), 139–140 (2010)

[14] Yan, F., Powell, D.R., Curtis, D.J., Wong, N.C.: From reads to insight: a hitchhiker’s guide to atac-seq data analysis. Genome biology 21, 1–16 (2020)

[15] Love, M.I., Huber, W., Anders, S.: Moderated estimation of fold change and dispersion for rna-seq data with deseq2. enome biology 15(12), 1–21 (2014)

[16] Lim, K.S., Reidenbach, A.G., Hua, B.K., Mason, J.W., Gerry, C.J., Clemons, P.A., Coley, C.W.: Machine learning on dna-encoded library count data using an uncertainty-aware probabilistic loss function. Journal of Chemical Information and Modeling (2022)

[17] Huisman, B.D., Dai, Z., Gifford, D.K., Birnbaum, M.E.: A highthroughput yeast display approach to profile pathogen proteomes for mhc-ii binding. eLife 11, 78589 (2022). https://doi.org/10.7554/eLife.78589

[18] Rappazzo, C.G., Huisman, B.D., Birnbaum, M.E.: Repertoire-scale determination of class ii mhc peptide binding via yeast display improves antigen prediction. Nature communications 11(1), 1–14 (2020)

[19] Bryant, D.H., Bashir, A., Sinai, S., Jain, N.K., Ogden, P.J., Riley, P.F., Church, G.M., Colwell, L.J., Kelsic, E.D.: Deep diversification of an aav capsid protein by machine learning. Nature Biotechnology 39(6), 691–696 (2021)

[20] Zhu, D., Brookes, D.H., Busia, A., Carneiro, A., Fannjiang, C., Popova, G., Shin, D., Chang, E.F., Nowakowski, T.J., Listgarten, J., et al.: Machine learning-based library design improves packaging and diversity of adeno-associated virus (aav) libraries. bioRxiv (2021)

[21] Wu, Z., Kan, S.J., Lewis, R.D., Wittmann, B.J., Arnold, F.H.: Machine learning-assisted directed protein evolution with combinatorial libraries. Proceedings of the National Academy of Sciences 116(18), 8852–8858 (2019)

[22] Boucher, J.I., Bolon, D.N., Tawfik, D.S.: Quantifying and understanding the fitness effects of protein mutations: Laboratory versus nature. Protein Science 25(7), 1219–1226 (2016)

[23] Bloom, J.D.: Software for the analysis and visualization of deep mutational scanning data. BMC bioinformatics 16(1), 1–13 (2015)

[24] Wijesooriya, K., Jadaan, S.A., Perera, K.L., Kaur, T., Ziemann, M.: Urgent need for consistent standards in functional enrichment analysis. PLoS computational biology 18(3), 1009935 (2022)

[25] Harvey, E.P., Shin, J.-E., Skiba, M.A., Nemeth, G.R., Hurley, J.D., Wellner, A., Shaw, A.Y., Miranda, V.G., Min, J.K., Liu, C.C., Marks, D.S., Kruse, A.C.: An in silico method to assess antibody fragment polyreactivity. bioRxiv (2022) https://www.biorxiv.org/content/early/2022/01/13/2022.01.12.476085.full.pdf.https://doi.org/10.1101/2022.01.12.476085

[26] Hu, D., Hu, S., Wan, W., Xu, M., Du, R., Zhao, W., Gao, X., Liu, J., Liu, H., Hong, J.: Effective optimization of antibody affinity by phage display integrated with high-throughput dna synthesis and sequencing technologies. PloS one 10(6), 0129125 (2015)

[27] Romero, P.A., Tran, T.M., Abate, A.R.: Dissecting enzyme function with microfluidic-based deep mutational scanning. Proceedings of the National Academy of Sciences 112(23), 7159–7164 (2015)

[28] Russ, W.P., Figliuzzi, M., Stocker, C., Barrat-Charlaix, P., Socolich, M., Kast, P., Hilvert, D., Monasson, R., Cocco, S., Weigt, M., et al.: An evolution-based model for designing chorismate mutase enzymes. Science 369(6502), 440–445 (2020)

[29] Poelwijk, F.J., Socolich, M., Ranganathan, R.: Learning the pattern of epistasis linking genotype and phenotype in a protein. Nature communications 10(1), 1–11 (2019)

[30] Song, H., Bremer, B.J., Hinds, E.C., Raskutti, G., Romero, P.A.: Inferring protein sequence-function relationships with large-scale positive-unlabeled learning. Cell systems 12(1), 92–101 (2021)

[31] Sarkisyan, K.S., Bolotin, D.A., Meer, M.V., Usmanova, D.R., Mishin, A.S., Sharonov, G.V., Ivankov, D.N., Bozhanova, N.G., Baranov, M.S., Soylemez, O., et al.: Local fitness landscape of the green fluorescent protein. Nature 533(7603), 397–401 (2016)

[32] Gelman, S., Fahlberg, S.A., Heinzelman, P., Romero, P.A., Gitter, A.: Neural networks to learn protein sequence–function relationships from deep mutational scanning data. Proceedings of the National Academy of Sciences 118(48), 2104878118 (2021)

[33] Segerman, B.: The most frequently used sequencing technologies and assembly methods in different time segments of the bacterial surveillance and refseq genome databases. Frontiers in Cellular and Infection Microbiology 10, 527102 (2020)

[34] Kanwar, N., Blanco, C., Chen, I.A., Seelig, B.: Pacbio sequencing output increased through uniform and directional fivefold concatenation. Scientific reports 11(1), 1–13 (2021)

[35] Rhoads, A., Au, K.F.: Pacbio sequencing and its applications. Genomics, proteomics & bioinformatics 13(5), 278–289 (2015)

[36] Gutmann, M., Hirayama, J.-i.: Bregman divergence as general framework to estimate unnormalized statistical models. arXiv preprint arXiv:1202.3727 (2012)

[37] Qin, J.: Inferences for case-control and semiparametric two-sample density ratio models. Biometrika 85(3), 619–630 (1998)

[38] Sugiyama, M., Suzuki, T., Kanamori, T.: Density-ratio matching under the bregman divergence: a unified framework of density-ratio estimation. Annals of the Institute of Statistical Mathematics 64(5), 1009–1044 (2012)

[39] Sugiyama, M., Suzuki, T., Kanamori, T.: Density Ratio Estimation in Machine Learning. Cambridge University Press, ??? (2012)

[40] Henaff, O.: Data-efficient image recognition with contrastive predictive coding. In: International Conference on Machine Learning, pp. 4182–4192 (2020). PMLR

[41] Oord, A.v.d., Li, Y., Vinyals, O.: Representation learning with contrastive predictive coding. arXiv preprint arXiv:1807.03748 (2018)

[42] Mohamed, S., Lakshminarayanan, B.: Learning in implicit generative models. arXiv preprint arXiv:1610.03483 (2016)

[43] Bartoli, L., Capriotti, E., Fariselli, P., Martelli, P.L., Casadio, R.: The pros and cons of predicting protein contact maps. Protein Structure Prediction, 199–217 (2008)

[44] Vendruscolo, M., Kussell, E., Domany, E.: Recovery of protein structure from contact maps. Folding and Design 2(5), 295–306 (1997)

[45] Brookes, D.H., Aghazadeh, A., Listgarten, J.: On the sparsity of fitness functions and implications for learning. Proceedings of the National Academy of Sciences 119(1), 2109649118 (2022)

[46] Perabo, L., Buüning, H., Kofler, D.M., Ried, M.U., Girod, A., Wendtner, C.M., Enssle, J., Hallek, M.: In vitro selection of viral vectors with modified tropism: the adeno-associated virus display. Molecular therapy 8(1), 151–157 (2003)

[47] Fox, E.J., Reid-Bayliss, K.S., Emond, M.J., Loeb, L.A.: Accuracy of next generation sequencing platforms. Next generation, sequencing & applications 1 (2014)

[48] Stöcker, B.K., Köster, J., Rahmann, S.: Simlord: simulation of long read data. Bioinformatics 32(17), 2704–2706 (2016)

[49] Reddi, S.J., Kale, S., Kumar, S.: On the convergence of adam and beyond. arXiv preprint arXiv:1904.09237 (2019)

[50] Newey, W.K., McFadden, D.: Large sample estimation and hypothesis testing. Handbook of econometrics 4, 2111–2245 (1994)

